# Discrete Community Assemblages Within Hypersaline Paleolake Sediments of Pilot Valley, Utah

**DOI:** 10.1101/634642

**Authors:** Kennda L. Lynch, Kevin A. Rey, Robin J. Bond, Jennifer F. Biddle, John R. Spear, Frank Rosenzweig, Junko Munakata-Marr

## Abstract

Hypersaline paleolake sediments are understudied ecosystems whose microbial ecology is largely unknown. Here we present mineralogical, geochemical, and small-subunit 16S rRNA gene sequence data on one such environment, the Pilot Valley Basin (PVB), a sub-basin of ancient Lake Bonneville located in northwest Utah. PVB exhibits a variety of aqueous minerals including phyllosilicates, carbonates, and sulfates, as well as microbially-induced sedimentary structures. As perchlorate occurs naturally (up to 6.5 ppb) in Pilot Valley sediments, and because recent evidence suggests that it is subject to biotic reduction, PVB has been proposed as a Mars analog site for astrobiological studies. 16S rRNA gene sequencing was used to investigate microbial diversity and community structure along horizontal and vertical transects within the upper basin sediments and beta diversity analyses indicate that the microbial communities in Pilot Valley are structured into three discrete groups. <u>O</u>perational taxonomic <u>u</u>nits (OTUs) belonging to the main archaeal phylum, Euryarchaeota, make up ~23% of the sequences, while OTUs belonging to three bacterial phyla, Proteobacteria, Bacteroides and Gemmatimonadetes, constitute ~60-70% of the sequences recovered at all sites. Diversity analyses indicate that the specific composition of each community correlates with sediment grain size, and with biogeochemical parameters such as nitrate and sulfate concentrations. Interestingly, OTUs belonging to the phylum Gemmatimonadetes are co-located with extreme halophilic archaeal and bacterial taxa, which suggests a potential new attribute, halophilicity, of this newly-recognized phylum. Altogether, results of this first comprehensive geomicrobial study of Pilot Valley reveal that basin sediments harbor a complex and diverse ecosystem.

## INTRODUCTION

Hypersaline ecosystems are globally distributed across a broad range of terrestrial and aquatic environments including saline flats, playas, soda lakes, saline lakes, hypersaline springs, solar salterns, and deep sea and oil-reservoir brines (Oren, 2006). Hypersaline ecosystems harbor diverse microbial communities often consisting of endemic taxa that represent all three domains of life (Andrei et al., 2012; Feazel et al., 2008; Oren, 2008; Robertson et al., 2009; Ley et al., 2006). To date, most microbial and biogeochemical studies of hypersaline environments have focused on aquatic habitats or subaqueous sediments whereas hypersaline soils and groundwater-dominated sediments have been largely neglected (Ventosa et al., 2008; Sirisena et al., 2018).

Hypersaline playas are often remnants of ancient lake basins, commonly known as paleolakes. Across the globe, numerous large paleolakes from the last ice age (chiefly freshwater/brackish lakes from the late Pleistocene/early Holocene Boundary) have gradually transitioned to modern-day hypersaline playas, e.g., the Chott el Gharsa of northern Africa, the Salar de Uyni of Bolivia, Death Valley in California, USA as well as the focus of this study, the Lake Bonneville Basin in northwestern Utah, USA (Barbieri & Stivaletta, 2012; Currey, 1990; Douglas, 2004; Fornari et al., 2001). During this transition, microbial life that initially dominated the water column and sediments would have been gradually replaced by halotolerant and halophilic microorganisms as water levels dropped and ions became more concentrated. Whether such changes occurred as a result of evolutionary adaptation by the original residents, by dispersal of novel taxa from other hypersaline environments, or by a combination of these processes is currently unknown. As brines in these systems became saturated, microbial activities may have accelerated the precipitation and/or production of minerals (i.e., biomineralization), and microbes may have become entrained in the resulting evaporites (Barbieri et al., 2006; Douglas & Yang, 2002). Because such basins tend to maintain closed groundwater systems that allow for continual wetting of the playa sediments, they continue to support modern-day microbial ecosystems (Hollister et al., 2010; Genderjahn et al., 2018; Sirisena et al., 2018; McGonigle et al., 2019). Surveys of microbial diversity in saline sediments indicate that they can be among the most taxonomically diverse communities known (Ventosa et al., 2008). However, little is known about the microbial ecology or the biogeochemical factors that drive community structure in hypersaline sedimentary environments, especially how microbes adapt to high salt concentrations under anoxic conditions (Schwendner et al., 2018).

Hypersaline paleolake basins have also been proposed as habitability analogs for similar environments on Mars, giving them astrobiological significance (Lynch et al., 2015). To date, the few studies that have evaluated microbial ecology in hypersaline sediments have done so within single vertical cores (Sirisena et al., 2018; Genderjahn et al., 2018). But closed basin systems exhibit horizonal as well as vertical geochemical and mineralogical gradients (Eugster & Jones, 1979), thus a study that examines microbial diversity in both dimensions is warranted. Here, we use multivariate statistics to integrate 16S rRNA gene sequencing with geochemical, mineralogical, and lithological analyses in order to discern which factors most strongly contribute to microbial distribution and abundance in Pilot Valley Basin, Utah. As our analyses encompass vertical and horizontal dimensions, we are able, for the first time, to make reasoned inferences about the relationship between microbial community structure and the geochemical and lithological characteristics of hypersaline lacustrine sediments. The information derived from studying this ecosystem could provide insight into the evolution and dynamics of hypersaline sediments elsewhere on Earth and beyond.

## MATERIALS AND METHODS

### Study site description

Ancient Lake Bonneville encompassed the majority of northern Utah: the modern-day remnants of the lake’s basin constitute the Great Salt Lake Desert (GSLD) and the Great Salt Lake (GSL). Bonneville was the largest of several North American freshwater paleolakes from the Pleistocene Epoch, covered 51,000 km^2^ of present-day western Utah and smaller sections of eastern Nevada and southern Idaho, and reached a maximum depth of ~300 meters. Around 14 ka., lake levels began a decline that eventually resulted in the modern playa system of the GSLD basin and the Great Salt Lake (DeRito & Madsen, 2008; Madsen et al., 2001; Spencer et al., 1984; Wilkerson, 2012). The modern GSLD, located in the Basin and Range physiographic province of North America, extends from the western edge of the Great Salt Lake into the eastern edge of Nevada, and is bifurcated into north/south sections by Interstate-80. It is classified as an arid desert with average temperatures ranging from 44°C in the summer to 7°C in the winter and an average annual precipitation of less than 150 mm/yr (Kottek et al., 2006; WRCC, 2013). Though it looks like a singular basin, the GSLD encompasses multiple enclosed sub-basins, largest of which are the Bonneville Salt Flats, the Pilot Valley basin and the Newfoundland basins (Jones et al., 2009), as shown in **Figure 1a**. Both the Newfoundland and Bonneville basins have undergone significant anthropogenic alteration, which has notably changed the brine chemistry and mineralogy of the Newfoundland Basin and has caused significant salt loss in the Bonneville basin (Jones et al., 2009; Mason & Kipp, 1997). The Pilot Valley basin (**Figure 1b**) has remained relatively isolated from anthropogenic influence, which adds to its strength as the chosen target for this study.

**Figure 1.**
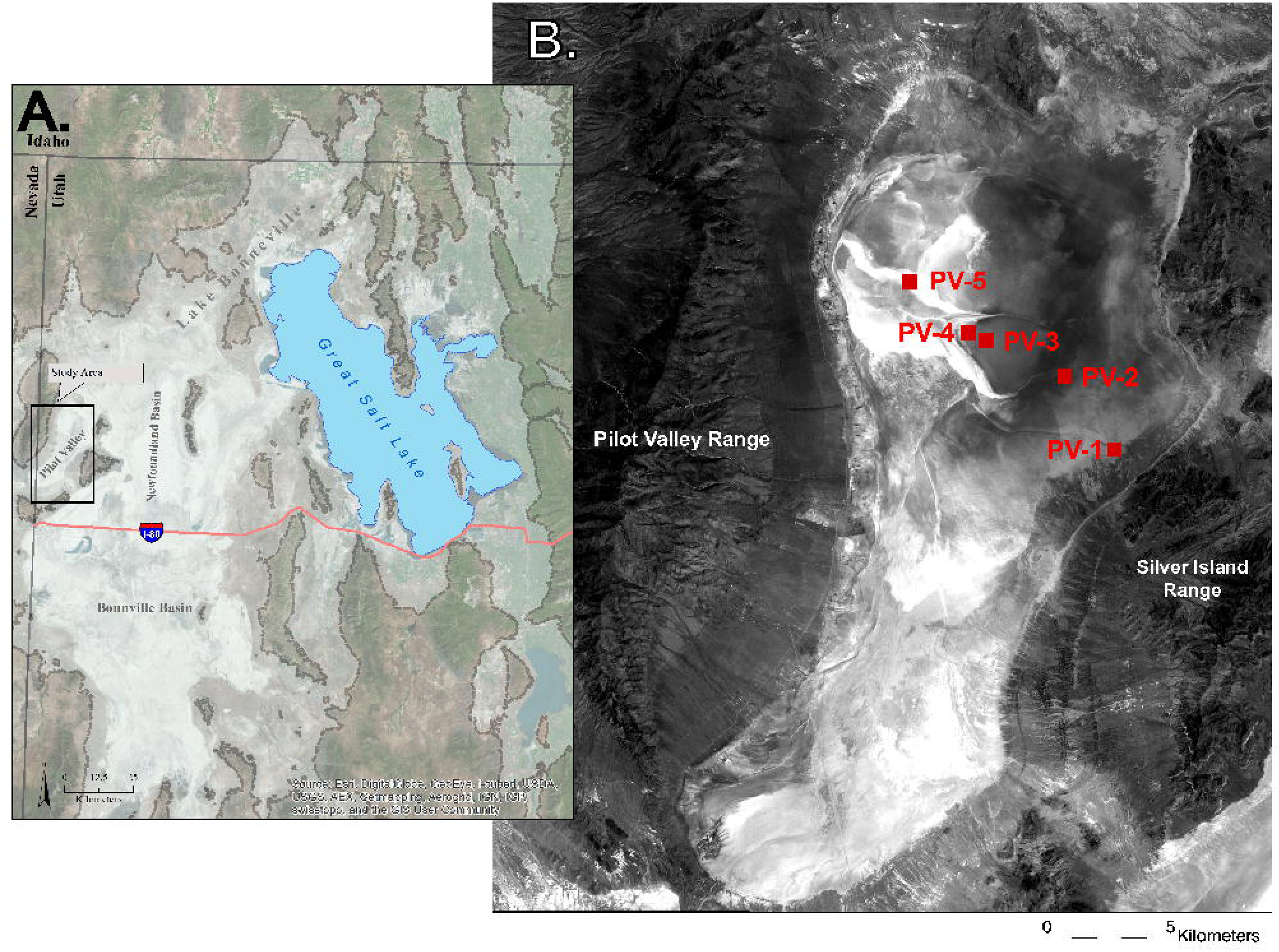
Field Site - Great Salt Lake Desert & Great Salt Lake. Inset - Pilot Valley Field site with sample site locations PV1 through PV5.

Pilot Valley is approximately 33 km in length and 8-15 km wide with the long axis running from northeast to southwest and the topographic center of the basin located in the northwestern corner (Lines, 1979). The basin exhibits classic closed-basin evaporite zonation due to differences in mineral solubilities that initially developed during the last regressive phase of Lake Bonneville, between 8,000 and 10,000 years ago (Eardley et al., 1957; Hunt, 1975; Lines, 1979). The least soluble minerals, mainly carbonates, were deposited around the rim of the basin, or were buried in the lower layers of the lacustrine sediments, in deposits that are described as a carbonate mud and occur either as a soft, puffy surface or a hard, compact surface. In the next zone, sulfates, specifically gypsum, intermix with the carbonate mud. Finally, within the hard “salt” pan at the topographic center of the basin, highly soluble chloride and Mg-sulfate salts are deposited. Extensive microbial mats are observed starting in the sulfate zone and propagating almost to the center of the basin. Due to continual reworking, mainly from episodic fluvial activity, many of the deposits are interbedded with mud layers originating from lower sedimentary deposits or clastic input (Lines, 1979).

Three main aquifers are associated with the basin: an alluvial fan aquifer, consisting of fresh to brackish water, present within the alluvial fans alongside each of the flanking mountain ranges; a deep basin-fill brine aquifer that underlies the entire basin at a depth of ~30 meters; and a shallow brine aquifer that encompasses the upper ~6 meters of the basin sediment fill (Mason & Kipp, 1997). This study focuses on fluids and sediments from the shallow brine aquifer, which is maintained by groundwater flow from mountain-front recharge of the alluvial aquifer flanking the Silver Island Range (Carling et al., 2012). It should be noted that hydrological connectivity between the deep basin-fill aquifer and the shallow brine aquifer is thought to be effectively non-existent (Mason & Kipp, 1997). Due to the frequency of recharge, Pilot Valley sediments remain consistently moist throughout the basin, although they are clearly more saturated at the center of the basin where the water table lies just below the surface. The only loss mechanism from the Pilot Valley basin is capillary wicking and evaporation (estimated rate 95 in/year) from the playa surface (Lines, 1979). Additional detailed field site characterization, including mineralogy, can be found in Lynch et al. (2015; 2019)

### Field sampling

Field samples were obtained on two sampling trips, June 2010 (**EX 1**) and May 2012 (**EX 4**). Sediment and aquifer fluid samples from Pilot Valley were collected along the same horizontal transect from the basin rim to the topographic center of the basin (**Figure 1**). *In situ* estimates of aquifer fluid pH ranged between 5.8-6.0; subsurface temperatures averaged ~20°C at the time of sampling for both expeditions. Subsurface temperatures were measured using an infrared thermometer and pH was measured using both pH strips and a portable pH meter. Sample cores were obtained using an AMS Extendible Core Sampler and recovery tripod, which retrieves 5 cm diameter cores up to 60 cm long. The cores were confined within plastic sleeves, which were cut using a hand-held circular saw to expose the core for inspection and sampling. DNA and geological samples were taken at each point where the lithology of the sediments clearly changed (**Figure S1**). Sediment core samples were collected and stored on CO_2_ ice in the field. Geological samples were stored at −20°C; samples for DNA extraction and analysis were stored at −80°C in the lab. Water from the basin aquifer was obtained from each bore hole (if possible) as well from permanent wells at the topographic center of the basin. Aquifer fluids were filtered to 0.2 µm, stored on H_2_O ice in the field then refrigerated at −4°C in the lab. Average free water content of the sediments was determined by weight. To estimate free water content, sediment sub-samples were defrosted to room temperature in the lab, weighed, oven-dried overnight at 90°C, then cooled and re-weighed.

### IC and ICP OES analysis

Pilot Valley fluid samples for ion chromatography (IC) analysis were diluted 10- and 100-fold prior to analysis; fluid samples for inductively coupled plasma-optical emission spectroscopy (ICP-OES) were also diluted 10- and 100-fold, then acidified with trace-metal grade nitric acid. All sediment samples were extracted for IC and ICP-OES following the Florida Department of Environmental Protection Method NU-044-3.13. Approximately 5 g of sediment were combined with 100 ml of deionized water in 250 ml Nalgene bottles. The bottles were shaken at 200 RPM for 30 minutes and allowed to settle overnight at 4°C. The resulting extracts were 0.2 µm filtered then diluted 10- and 100-fold; extracts for ICP-OES analysis were also acidified with trace-metal grade nitric acid. All IC samples were analyzed for major anions using a Dionex ICS-90 ion chromatography system with an AS14A (4x 250 mm) column at the Colorado School of Mines. ICP-OES samples were analyzed for major cations using a Perkin-Elmer Optima 5300 DV spectrometer at Mines. Mineralogical analysis of each sample site was published in Lynch et al. (2015).

### Total carbon analysis

Sediment samples were weighed, dried overnight at 100°C, then ground with a mortar and pestle in preparation for carbon analysis. Total inorganic (TIC), total organic (TOC), and total carbon (TC) analyses were conducted using a UIC Carbon Analyzer System (Joliet, IL) at Mines. Total inorganic carbon was determined using a UIC CM5130 Acidification Module. After placing approximately 10 mg of sample in a heated glass vial, 2 ml of 2% H_2_SO_4_ were added to dissolve inorganic carbon, which was then released as CO_2_ gas. The effluent was collected and quantified by a UIC CM5014 Coulometer for total inorganic carbon by % weight. Total carbon was determined using a UIC CM5300 Furnace Module. Approximately 20-40 mg of sample was placed in a ceramic crucible, heated to ~935°C where all organic and inorganic carbon was incinerated to release CO_2_ gas. The effluent was also collected and quantified by a UIC CM5014 Coulometer for total carbon by % weight. Both TIC and TC were repeated in triplicate to ensure a measurement error less than 10%. TOC was calculated as the difference in TC and TIC.

### DNA extraction, PCR, qPCR and DNA sequencing

Bulk DNA was extracted using the PowerSoil DNA Isolation Kit (Qiagen, Catalog # 12888; formerly MoBio Laboratories). DNA was extracted from ~25 mg of each sample in triplicate, following the manufacturer’s protocol with two modifications to maximize DNA recovery, concentration, and purification. The first modification was at the clean-up step, as the Pilot Valley samples contain compounds (e.g., heavy metals, humics, etc.) that can inhibit the <u>p</u>olymerase <u>c</u>hain reaction (PCR). Hence, the C3 solution humic removal step (MoBio Powersoil DNA Isolation Kit Instruction Manual (https://www.qiagen.com)) was repeated twice to ensure DNA purity. The second modification was the last step where DNA from each replicate was concentrated onto the same single spin filter, eluted into a single tube and stored at −20°C. In preparation for 16S rRNA gene sequencing, DNA was PCR amplified and barcoded using the small sub-unit (SSU) rRNA gene primers 515f-modified forward primer, 5’- GTGYCAGCMGCCGCGGTAA-3’ that also included an 8-base barcode and a 927r-modified reverse primer, 5’-CCGYCAATTCMTTTRAGTTT-3’, which covers the V4 and V5 region of the 16S rRNA gene, as per Osburn et al. (2011). Quantitative PCR (qPCR) amplification was carried out on a LightCycler 480 II (Roche Applied Sciences) using touchdown PCR and terminated after reaching the plateau stage in order to minimize primer dimer formation. Samples were amplified in 21 μL reactions, each of which contained, per reaction: Phusion Master Mix (New England BioLabs, Inc.), 3% final volume DMSO, 0.4X final concentration SYBR Green (stock concentration was 40X), 500 nM final concentration of each primer, and 2 μl of template DNA. Final amplicon concentration was determined using an Agilent Bioanalyzer 2100 (Agilent Technologies). Samples were gel-purified (Montage DNA Gel Extraction Kit, Millipore) then normalized and pooled to achieve an equal concentration of amplicon DNA from each sample. The pooled amplicons were vacuum evaporated to a desired final volume and shipped to Engencore (Columbia, SC) for sequencing on the 454-titanium pyrosequencing platform (Roche Applied Sciences).

### Processing, quality control and statistical analysis of 16S rRNA

The resulting partial SSU rRNA sequences were processed in QIIME 1.8 (Caporaso et al., 2010). Sequences and barcodes were de-multiplexed using the split_libraries.py script with the default parameters. Sequences were then de-noised using the low-level interface to the QIIME de-noiser. Sequences were chimera checked using USEARCH 6.1 then clustered into <u>o</u>perational taxonomic <u>u</u>nits (OTUs) using UCLUST at 97% similarity. OTUs were picked *de novo* using UCLUST with default parameters. The OTU representative sequences were aligned using PyNAST and the Greengenes 13_8 aligned reference database. Taxonomy was assigned using UCLUST and the Greengenes 13_8 97% similarity taxonomy reference database. Statistical analyses were conducted using QIIME and PAST software packages (Caporaso et al., 2010; Hammer et al., 2001). All DNA sequence data related to this study can be obtained through the European Nucleotide Archive (ENA) via accession number PRJEB11779.

## RESULTS

A total of 28 samples was collected for microbial community analysis (**Table 1**). For basin comparison analysis, samples were collected from the Pilot Valley basin rim, from the Bonneville Salt Flats (BSF) basin rim, and from exposed lake sediments from the rim of the northern arm of the GSL. For the detailed transect study of Pilot Valley, four sample cores were collected and subsampled along a horizontal geochemical gradient from the Pilot Valley basin rim to the topographical center (**Figure 1** inset, **Table 1**).

**Table 1.** Sample Identification & Depth

| Site ID | Core ID | Depth (cm) | Expedition |
| --- | --- | --- | --- |
| GSL-SJ | N/A | 0-45 | EX1 |
| BSF-Rim | N/A | 0-45 | EX1 |
| PV-Rim | N/A | 0-45 | EX1 |
| PV-1 | 1 | 0-3 | EX4 |
|  | 2 | 3-5 | EX4 |
|  | 3 | 8-10 | EX4 |
|  | 4 | 10-48 | EX4 |
|  | 5 | 48-51 | EX4 |
| PV-2 | 1 | 0-13 | EX4 |
|  | 2 | 13-22 | EX4 |
|  | 3 | 22-29 | EX4 |
|  | 4 | 29-42 | EX4 |
|  | 5 | 61-71 | EX4 |
|  | 6 | 71-94 | EX4 |
|  | 7 | 94-107 | EX4 |
|  | 8 | 107-117 | EX4 |
|  | 9 | 122-130 | EX4 |
|  | 10 | 130-152 | EX4 |
|  | 11 | 152-168 | EX4 |
| PV-3 | 1 | 0-8 | EX4 |
|  | 2 | 8-17 | EX4 |
|  | 3 | 17-32 | EX4 |
| PV-5 | 1 | 0-1 | EX4 |
|  | 2 | 1-6 | EX4 |
|  | 3 | 6-12 | EX4 |
|  | 4 | 12-28 | EX4 |

### Pilot Valley geochemistry and sedimentology

In addition to the mineralogical composition described in the literature (Lines, 1979; Lynch et al., 2015; Rey et al., 2016), both the Pilot Valley (PV) sediments and aquifer brines host an array of mobile constituents, with Cl, Na, SO_4_, K, Ca, and Mg present in the highest concentrations across the transect (**Tables S2, S3)**. The ion concentrations and pH define Pilot Valley as an athalassohaline-type hypersaline system (Ventosa & Arahal, 2009). Consistent with previous reports (Carling et al., 2012; Lines, 1979; Lynch et al., 2015; Warren, 1999; Rey et al., 2016), the average free water content across the basin is 27% ±3.6 (**Table S4**), indicating that the aquifer fluids encompass most of the subsurface environment. In addition to water content, certain trends can immediately be seen across the transect. For example, compared to the outer basin at PV1, soluble Ca and SO_4_ are more abundant in the inner basin sediments at PV3 & PV5 and at the transition zone, PV2 (one-way ANOVA P ≤ 0.01; Kruskal-Wallis P ≤ 0.05) (**Table S2**). These results correlate with previous mineralogical analyses of the basin and confirm the endorheic nature of Pilot Valley basin (Lynch et al., 2015). During the final regressive phases of Lake Bonneville, carbonates would have deposited in the outer basin and the majority of sulfates would have begun to deposit in the transition zone near PV2, with continued deposition into the topographic center of the basin (Lines, 1979; Lynch et al., 2015). Thus, quantitative measurements of geochemical, carbon, and water content can all be linked to the depositional history of Lake Bonneville.

Physical characteristics such as grain size are also linked to this depositional history. The depositional history of Lake Bonneville has been extensively studied in various parts of the basin over the past 100 years, which has resulted in a general overarching depositional timeline for all of Lake Bonneville, including Pilot Valley (Oviatt, 2015; Rey et al., 2016). Stratigraphic analysis of the upper 5 meters of sediment in Pilot Valley was conducted by Rey et al. (2016), resulting in five stratigraphic units representing sediment deposition in Pilot Valley that can be correlated to the depositional history of Lake Bonneville. Based on Rey et al.’s analysis, the sediments in our study represent two of these five stratigraphic units: Unit I and Unit II. Unit I is identified as laminated silty and sandy sediments, and represents the last regressive phase of Lake Bonneville to the present-day playa. Unit II is identified as a transition from an olive gray marl to a yellow gray marl and represents the transition in lake levels most likely from the Bonneville to Provo levels. It should be noted that the extensive salt deposit found in the north-central salt pan (also the topographic center of the basin) is independent of the defined units, as it formed after Lake Bonneville was gone. Secondary features also developed over time within the basin sediments after the main lake waters had regressed. These secondary features include *in situ* precipitated salt grains that over time created discontinuous beds of salt crystals, which then created secondary porosity that allowed for increased fluid flow throughout the basin sediments.

We determined grain size of the sedimentary layers qualitatively, based on comparing core images collected for this study in relation to previous descriptions of Pilot Valley stratigraphy and lithology as described above. Based on this visual analysis and identification of these sedimentary features in the cores, three grain size classifications were identified that were consistent with material seen in Units I and II (Figure 2, Table 2): fine clay (< 3 μm), fine silt (3 μm to 500 μm), and coarse grains (> 500 μm). The fine clay grain type is mainly a combination of illite, kaolinite and chlorite clays mixed with carbonate mud. The fine silt is mainly a carbonate mud composed of calcite, aragonite, dolomite and other carbonates, mixed with small sulfate and chloride salts, and some minor phyllosilicate concentrations. The coarse grains are large euhedral salt crystals, most likely calcium sulfate (gypsum) mixed with calcium chloride (halite) and other minor salts. When the Pilot Valley geochemistry of the sub-cores (**Tables S2-S4**) is rearranged to consider the three grain size classifications, some very clear geochemical trends emerge that are discussed below (**Tables 3**, **Tables S8-S10**).

**Figure 2.**
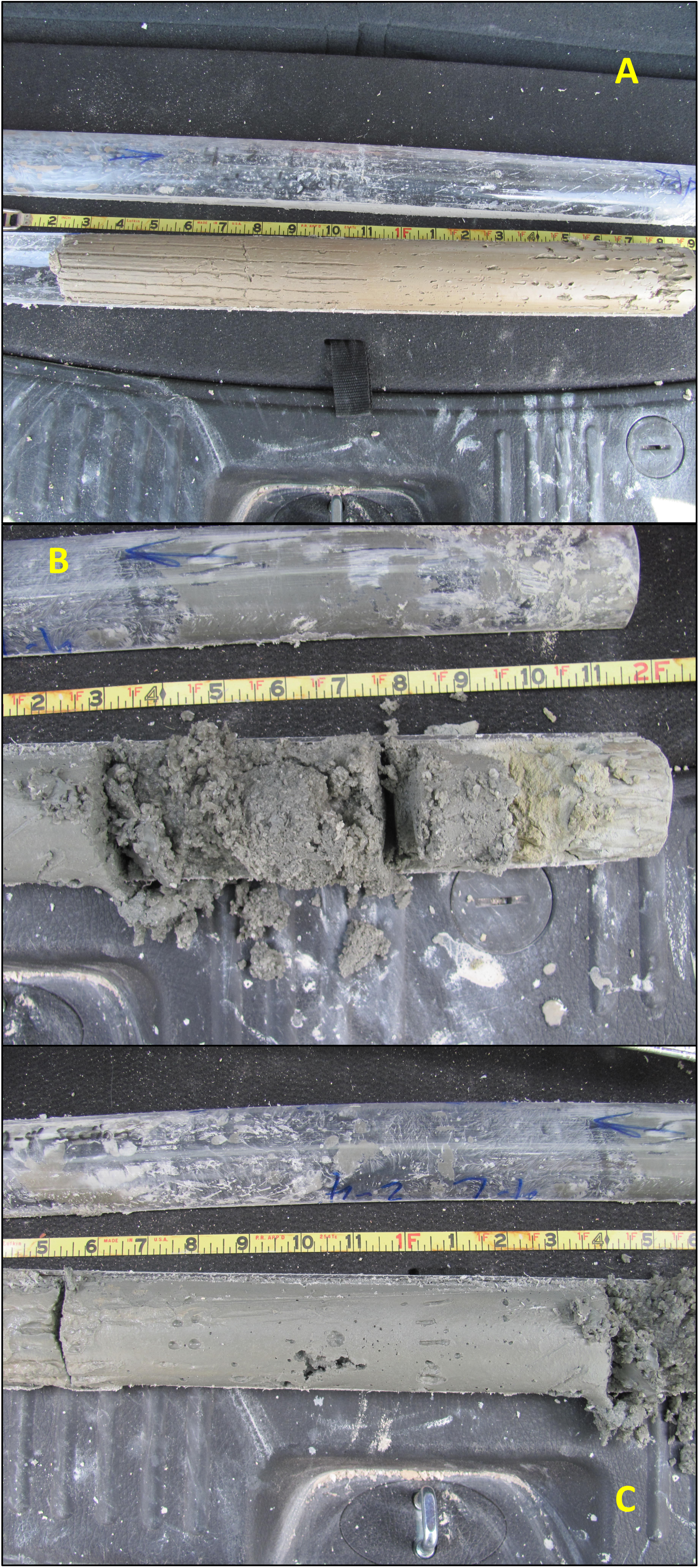
Images of the three grain types: A. Fine Clay, B. Fine Silt, C. Coarse

**Table 2.** Cluster Groups

| Table 2. Cluster Groups |  |
| --- | --- |
| ***Group 1*** |  |
| PV1-1, PV1-2, PV2-2,<br>PV2-3, PV2-4, PV2-5,<br>PV5-4 | Grain Type - Fine Clay |
|  | Clearly seen fine compacted clay sized particles. (Figure S8a) |
| ***Group 2*** |  |
| PV2-6, PV2-7, PV2-8,<br>PV2-9, PV3-2, PV3-3 | Grain Type - Coarse |
|  | Clear large euhedral salt crystals. (Figure S8c) |
| ***Group 3*** |  |
| PV1-3, PV1-4, PV1-5,<br>PV2-1, PV2-10, PV2-11,<br>PV3-1, PV5-1, PV5-2, PV5-3 | Grain Type - Fine Silt |
|  | Silt/sand-sized particles with some small evaporite crystals in the mix. (Figure S8b) |

### Microbial diversity of Pilot Valley compared to other Bonneville Basin remnants

Relative to aquatic environments, hypersaline soils and sediments like those in the Bonneville Basin have been neglected by microbiologists. For example, while studies of the microbial ecology of the Great Salt Lake began with Post (1977) and continue today, microbial diversity in other Bonneville Basin remnants remains largely unknown (Kjeldsen et al., 2007; Tazi et al., 2014; Lindsay et al., 2017; Baxter et al., 2005; Meuser et al., 2013; Boyd et al., 2017). Several studies have been conducted on the BSF, albeit focused on specific metabolisms of interest or solely on the archaeal domain (Myers & King, 2017; Boogaerts, 2015). Further, previous geological studies suggested that the mineralogical composition of the BSF and Pilot Valley were the same (Turk et al., 1973; Lines, 1979), implying that the microbial ecology might be similar. However, our recent geochemical characterization of the basin shows that the mineral composition of the two regions differ (Lynch et al., 2015). Given past geological work on both basins, we first compared microbial diversity of basin sediments in relation to one another and to that in a well-known standard: the GSL. Samples were therefore collected from the rims of Pilot Valley, the BSF, and the northern (hypersaline) arm of the GSL (**Figure 1**).

Samples sequenced from these sites yielded, after processing and quality control, a total library of 55,754 high-quality sequences (**Table S1**). Sequences were clustered into 1769 OTUs having 97% similarity. Five percent, 1%, and 10% of sequences were unassigned in Pilot Valley, the BSF and the GSL, respectively. In the archaeal domain across all three sample sites, a majority of sequences (≥99%) mapped to the phylum Euryarchaeota and the class/order/family *Halobacteria/Halobacteriales/Halobacteriaceae*. The only differentiation occurred at the genus level, with > 30% of Pilot Valley sequences mapping to an unknown genus, the BSF sequences mapping chiefly to the genus *Halomina*, and the GSL sequences mapping chiefly to the genus *Haloarcula*. Altogether these data suggest limited alpha and beta diversity within the archaeal domain. However, considerably more site-specific sequence variation was observed within the bacterial domain.

For Pilot Valley, 34% of the sequences mapped to the phylum Gemmatimonadaetes, 30% to Bacteroidetes, and 11% to Proteobacteria (of which 93% were *Gammaproteobacteria* and 7% *Deltaproteobacteria*) (**Figure 3**). By contrast, for both the BSF and the GSL, the majority of sequences mapped to Bacteroidetes phyla (65% and 58%, respectively), and specifically to the genus *Salinibacter* (38% and 51%, respectively). The second largest group of BSF and GSL sequences mapped to Proteobacteria (19% and 34% respectively), primarily *Deltaproteobacteria* (18% and 34%, respectively), which includes many <u>s</u>ulfate-reducing <u>b</u>acteria (SRB). The prevalence of Deltaproteobacteria in BSF and GSL contrasts with the Pilot Valley sample, where less than 1% of SSU sequences were *Deltaproteobacteria*. <u>P</u>rincipal <u>co</u>ordinate <u>a</u>nalysis (PCoA) of Bray-Curtis, weighted, and unweighted Unifrac metrics (**Figure S3**) show that the microbial communities in GSL and BSF sediments are more taxonomically similar to each other than they are to Pilot Valley sediments. Especially noteworthy in PV sediments, relative to these other sites, are the abundance of OTUs belonging to the phylum Gemmatimonadetes and the scarcity of OTUs belonging to the Deltaproteobacteria (< 1%), which include many sulfate reducing bacteria.

**Figure 3.**
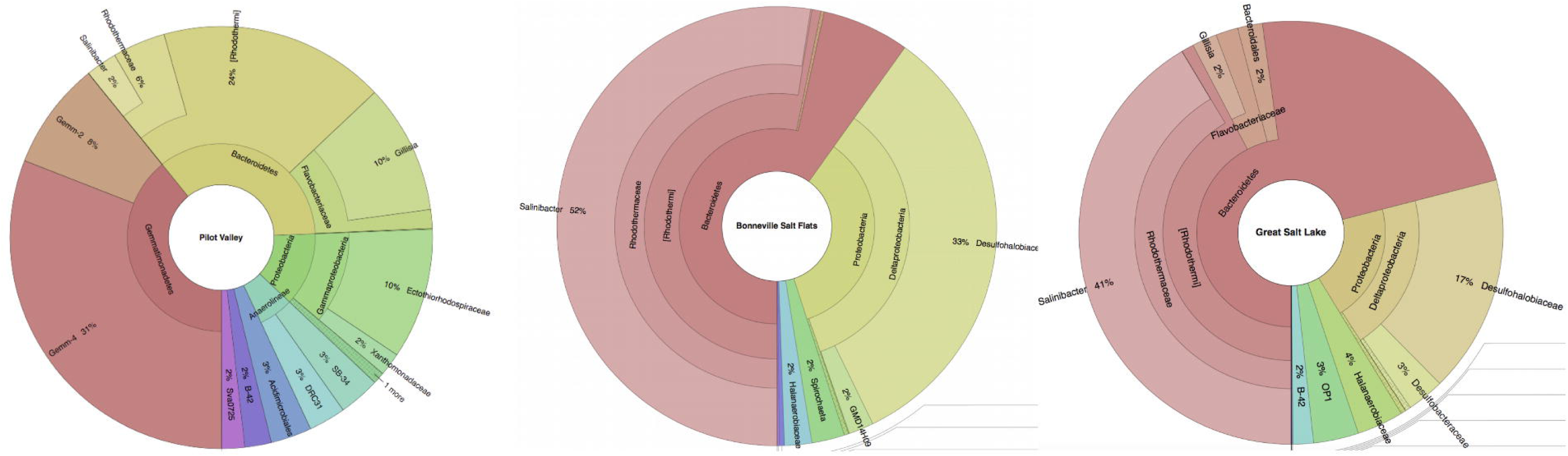
Krona plots of comparison microbial diversity at Pilot Valley, Bonneville Salt Flats, and the northern arm of the Great Salt Lake.

### Community composition along a horizontal transect in Pilot Valley

Samples sequenced from the Pilot Valley transect yielded a total library of 190,806 high quality sequences (i.e., Q scores of 30 or above) post processing and quality control (**Table S1**). These sequences were clustered into 1336 OTUs each having > 97% similarity; rarefaction analysis indicated that the sequencing effort provided reasonable coverage of diversity across the transect (**Figure S4**).

Archaeal sequences within the Pilot Valley transect are dominated by Halobacteria (99%) across all four sample locations. The majority of bacterial sequences across the Pilot Valley horizontal transect mapped to the phyla Proteobacteria (PV1=37%, PV2=50%, PV3=60%, PV5=58%), Gemmatimonadetes (PV1=34%, PV2=13%, PV3=4%, PV5=15%), and Bacteroidetes (PV1=13%, PV2=22%, PV3=19%, PV5=4%), though in PV3 the candidate phylum Acetothermia made up 6% of the population whereas in PV5 Acidobacteria made up 9% of the population (**Figure 4**). Almost no SRB-containing taxa from the Deltaproteobacteria (<1%) were seen at PV1 and PV5. However, known SRB-containing taxa were observed at PV2 (15%) and at PV3 (12%). It should also be noted that while cyanobacteria are visibly evident in microbial mats dispersed throughout the surface of Pilot Valley basin between PV-2 and PV-5, the relative abundance of cyanobacterial sequences in our dataset is small. This is likely due to the thin morphology and discontinuous distribution of these mats, though we cannot rule out the possibility of DNA extraction bias (Couradeau et al., 2011; Morin et al., 2010). Beta diversity analysis of depth-averaged community structure along the horizontal transect indicates that overall, despite the alpha diversity, populations between sample sites did not significantly differ. This observation suggests minimal phylogenetic variability along the horizontal evaporative gradient, even though taxonomic abundance varies greatly (Bray-Curtis R^2^=0.16, P_ADONIS_=0.27; weighted Unifrac R^2^=0.13, P_ADONIS_=0.45). The diversity of the vertical transect exhibited a much higher degree of phylogenetic variability (below).

**Figure 4.**
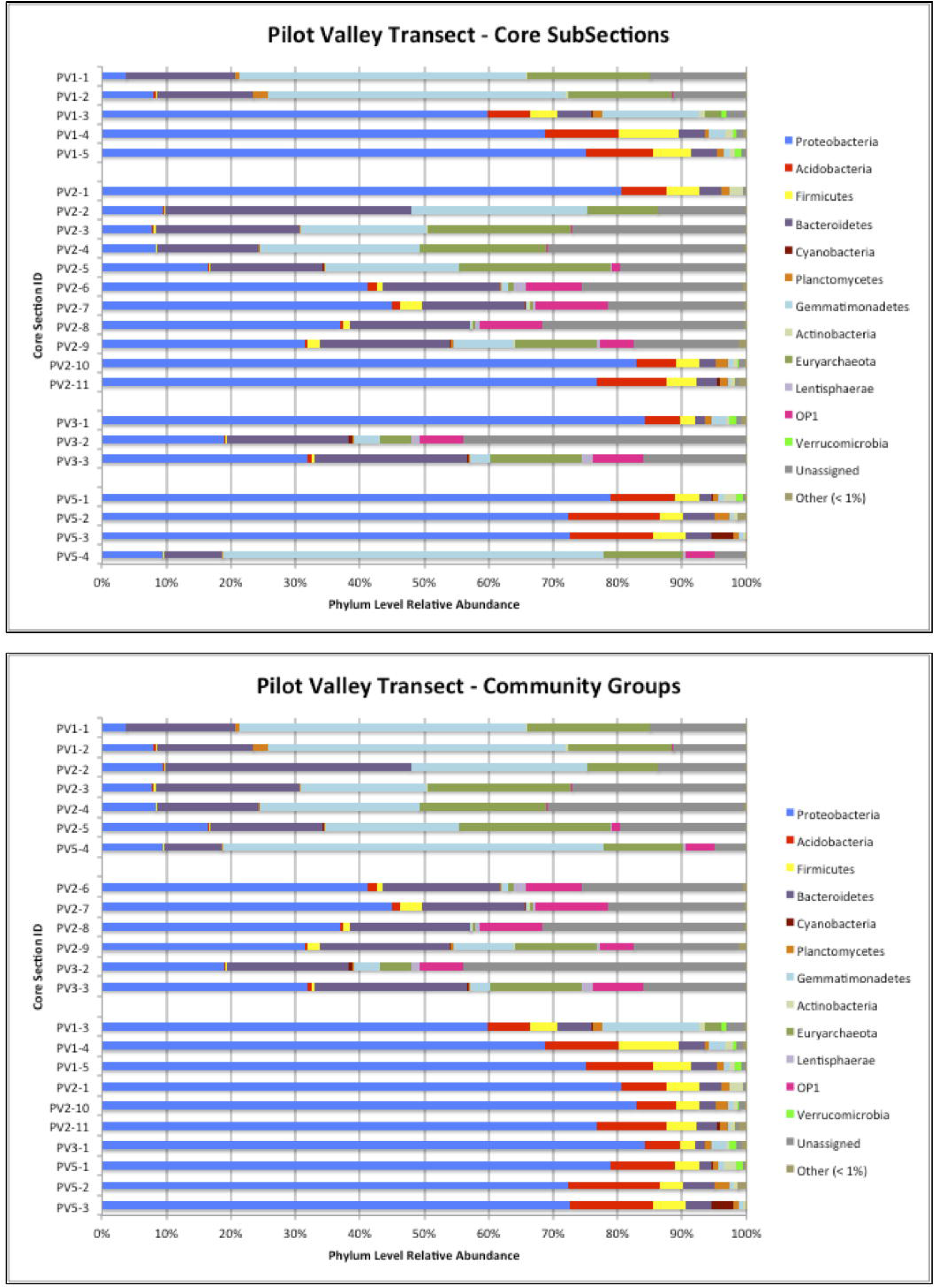
Phylum level relative abundances for horizontal transect study.

### Community composition within vertical transects in Pilot Valley

At all four Pilot Valley sites, the structure of microbial communities shifts abruptly between discrete layers within vertical transects down to a depth of ~2 m (**Figure 5a**), especially with regard to the phyla most abundantly represented. PCoA and PERMANOVA analysis of the Bray-Curtis (R^2^=0.46, P_ADONIS_=0.001), Morisita-Horn (R^2^= 0.52, P_ADONIS_=0.001), unweighted Unifrac (R^2^= 0.25, P_ADONIS_=0.001) and weighted Unifrac (R^2^= 0.61, P_ADONIS_=0.001) metrics show that sub-core samples cluster into three distinct groups; only in the unweighted Unifrac measure do samples PV5-4 and PV2-5 cross from Group 1 to Group 2 (**Figure 6**, **Table 2**, **Figure 5b**). These three groups also correlate with grain size classifications illustrated in **Table 2**: Group 1 correlates with fine clay; Group 2 with coarse grains; and Group 3 with fine silt. Remarkably, within these three groups, dominant taxonomic patterns are evident down to the genus level.

**Figure 5.**
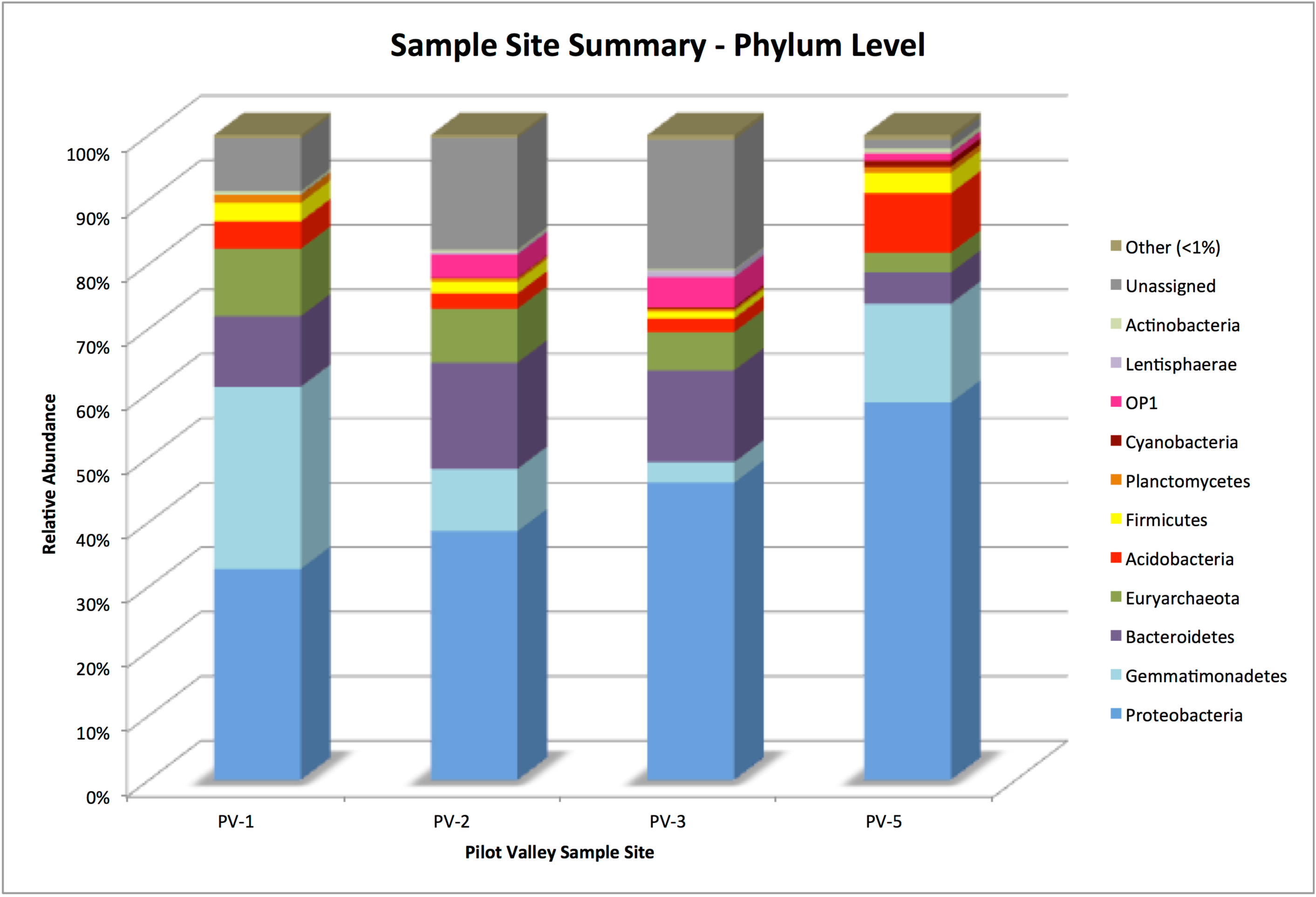
Phylum level relative abundances for vertical transect study. **A.** Core sub-sections, **B.** Community groups

**Figure 6.**
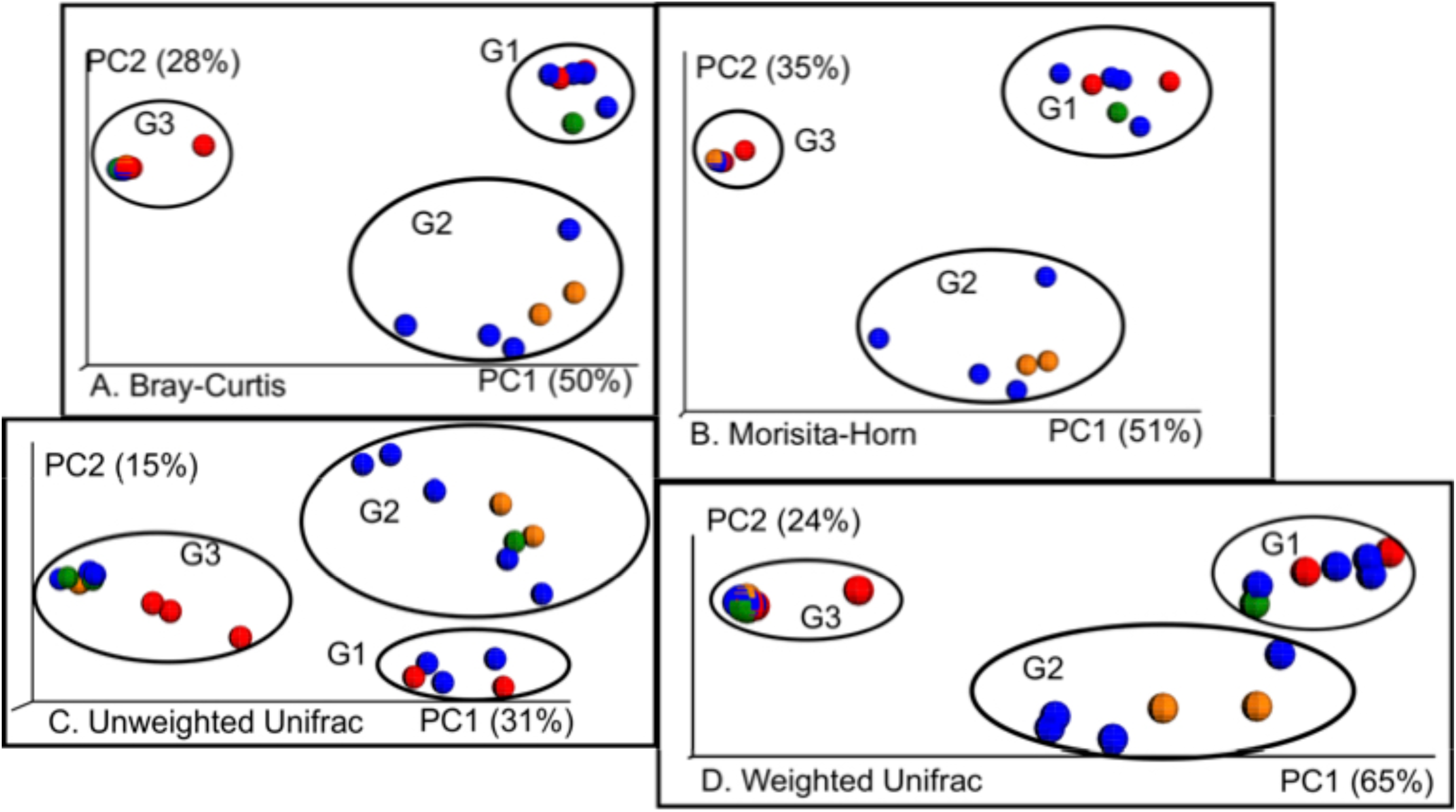
PCoA Plots of **A.** Bray-Curtis, **B.** Morisita Horn, **C.** Unweighted Unifrac, and **D.** Weighted Unifrac beta diversity metrics on core subsamples across Pilot Valley Transect. Sample Sites are identified by color: Red = PV-1, Blue = PV-2, Orange = PV-3, Green = PV-5.

In Group 1 (fine clay), the dominant taxa are order/family *Halobacteriales/Halobacteriaceae* (20%) phylum/class Gemmatimonadetes*/Gemm-4* (44%), genus *Salinibacter* (20%), and class Gammaproteobacteria (7%). Within the Group 1 *Gammaproteobacteria*, the dominant clade is the genus *Halomonas* at 50% (4% of total bacteria). In Group 2 (coarse grains), the dominant taxa are phylum Bacteroidetes (35%), class *Gammaproteobacteria* (15%), phylum Acetothermia (16%), and order *Halobacteriales* (9%). Gemmatimonadetes are only 6% of total assigned sequences in Group 2, and within the Bacteroidetes, only 15% (5% of total assigned sequences) are genus *Salinibacter*. Within the G*ammaproteobacteria*, 63% (12% of total assigned sequences) are genus *Halomonas*. In Group 3 (fine silt), the dominant taxa are genus *Halomonas* (54%), genus *Shewanella* (12%), and phylum Acidobacteria (8%). Beta diversity analyses indicate that Group 1 and Group 2 are more similar to one another than to Group 3. Unifrac metrics help delineate the nature of this relationship as the unweighted metric suggests a close relatedness between taxa present in both Groups 1 and 2. However, the difference between these two groups in weighted Unifrac suggests an environmental influence that enables different taxa to thrive in different regions (Lozupone et al., 2007).

Mantel correlation results for both weighted and unweighted Unifrac metrics (**Table S7**) show that there is little to no correlation between community structure and most of environmental variables measured (though Mantel correlations may not be as powerful as other multivariate methods (Legendre & Fortin, 2010)). While calcium, nitrate and sulfate all exhibit significant positive correlation to the community in the unweighted Unifrac metric, the weighted Unifrac shows almost no correlation between community structure exists and these environmental parameters. This suggests that while relative abundance of community members is not correlated to these parameters, community structure and phylogenetic relatedness is. Only barium (discussed below) is significant across both metrics, and it is weakly positive (r < 0.05).

Taxonomic data within Pilot Valley cores were constrained in a <u>c</u>anonical <u>c</u>orrespondence <u>a</u>nalysis (CCA) by eight of the best correlated quantitative environmental variables and one qualitative variable (grain size); this procedure resulted in an optimal eigenvalue for axis 1 of 0.428 (explaining 66.1% of the inertia; *P*=0.04) and an eigenvalue for axis 2 of 0.20 (explaining 22.48% of the inertia; *P*=0.04) (**Figure 7**). CCA plots confirm that the communities organize into the same three distinct Groups identified by PCoA (**Figure 6**, **Figure 7a**, **Table 2**), and show which environmental variables are best correlated with each group. Group 1 is associated with high barium and nitrate concentrations, and is also qualitatively related to fine clay-sized particles. Group 2 is mostly related to high total organic carbon (TOC), strontium, sulfate and calcium concentrations, and is linked to coarse grains and correlated with low nitrate. Group 3 is tightly clustered and correlated with increased water content, low nitrate, and fine silt grains.

**Figure 7.**
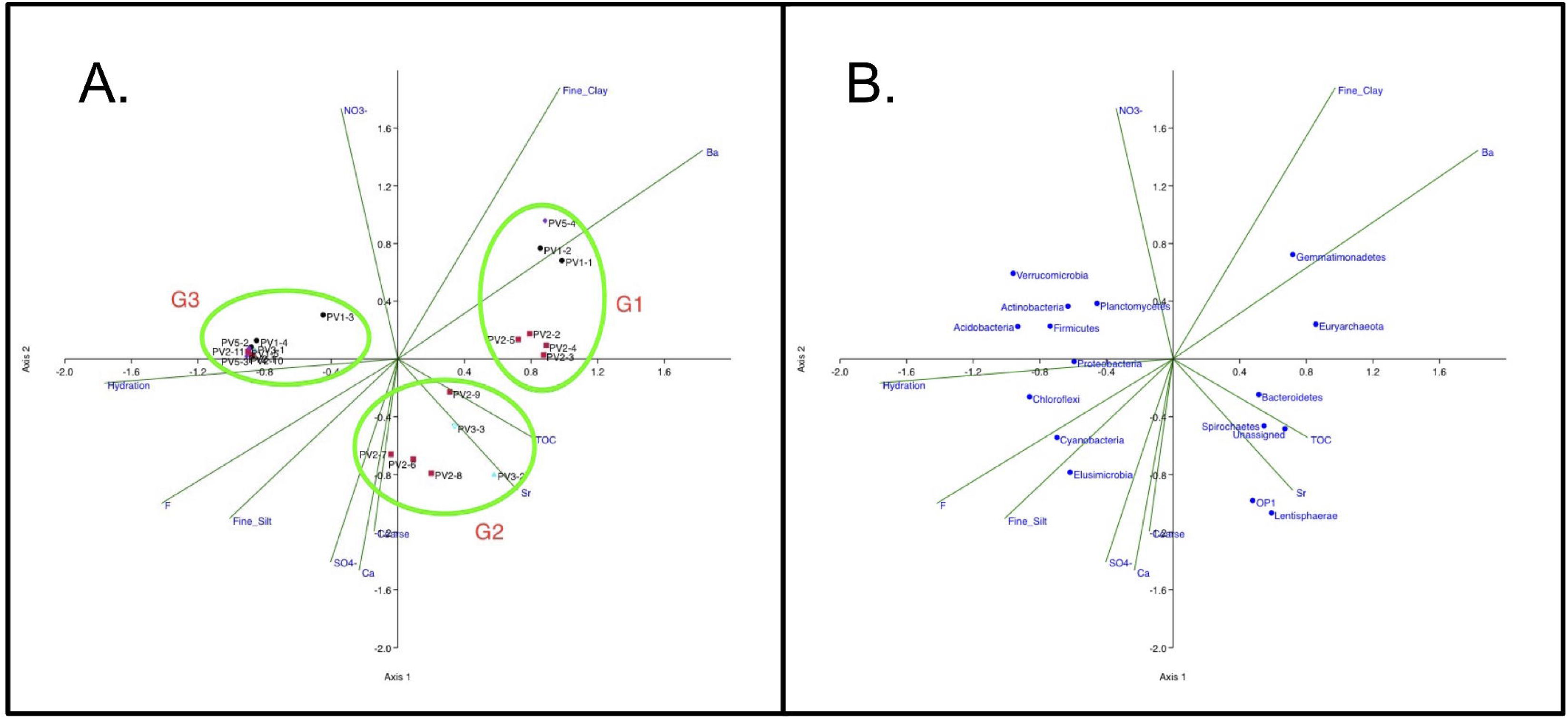
**A.** CCA plot of vertical cores subsections constrained by Pilot Valley environmental factors - scale factor 1. Groups identified as G1-Group1, G2-Group 2, G3-Group 3. Group 3 Members: PV1-3, PV1-4, PV1-5, PV2-1, PV2-10, PV2-11, PV3-1, PV5-1, PV5-2, PV5-3. **B.** CCA plot of phylum-level group distributions with respect to environmental constraints - scale factor 2.

Both Gemmatimonadetes and Euryarchaeota, the dominant phyla in Group 1 (fine clay), are correlated with high barium concentrations and clay-sized sediments (**Figure 7b**), as are most archaea. Bacteroidetes, a dominant phylum in Group 1 and in Group 2 (coarse grains), is correlated with increasing strontium concentrations and also increasing TOC concentrations as are Spirochaetes, Acetothermia, Lentisphaerae, and the unassigned sequences. Elevated strontium likely reflects increased salinity in both the Group 1 and Group 2 sediment types, as this element is known to be concentrated in brine waters and in sediments where there is salt mineral dissolution (Kloppmann et al., 2001). The strong correlation of Acetothermia (OP1) with increasing TOC is a reasonable result as this phylum contains acetogens. The strong correlation of TOC with Lentisphaerae is also to be expected as taxa within this phylum are known to produce <u>e</u>xtracellular <u>p</u>olymeric <u>s</u>ubstances (EPS) that would result in an increased carbon signature (Choi et al., 2015).

All remaining phyla are correlated with high water content. The primary cluster containing Firmicutes, Actinobacteria, Acidobacteria, Verrucomicrobia, and Planctomycetes correlates strongly with increasing hydration and increasing nitrate. This finding is consistent with the fact that each of these groups host taxa that carry out parts of the nitrogen cycle (Delmont et al., 2018; Mohammadi et al., 2017; Gtari et al., 2012; Ward et al., 2009; Heylen & Keltjens, 2012). Also, when the geochemistry of all the samples is reorganized by community group (Table 3, Table S9) we see clear evidence for nitrate utilization in the Group 2 sediments as nitrate concentrations from all samples fall below the limit of detection. Proteobacteria, Chloroflexi, Cyanobacteria, and Elusibacteria are all also associated with increasing hydration, fine silt, and elevated fluorine concentrations. The fluorine data suggests increased concentrations of carbonates (calcite, aragonite, and dolomite) as they are known to contain high fluorine content (Rude & Aller, 1991).

**Table 3.**
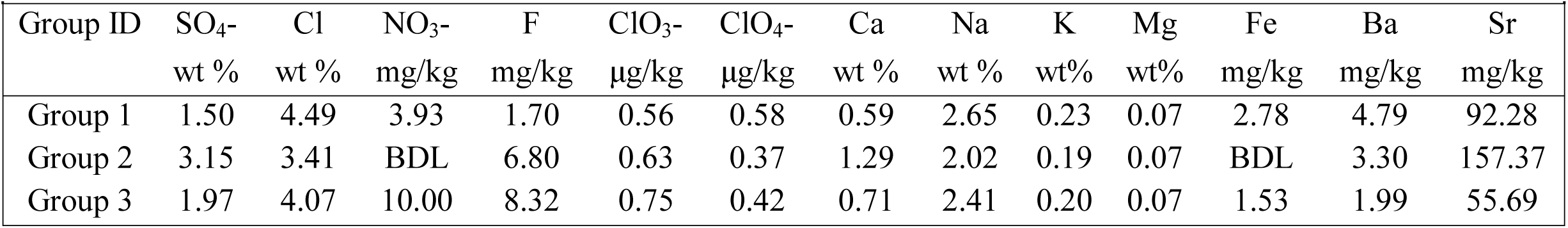
Pilot Valley Geochemistry - Sediment Ions by Group

## DISCUSSION

### Community assemblages correlate with grain size classification

Altogether, twenty-two environmental variables were measured at Pilot Valley (**Tables S2-S5** and **Table 2**). Of those variables, the sediment grain size classification appears to be the one most highly correlated with how microbial communities assemble. Grain size has previously been evaluated as contributing to community assemblages in aquatic benthic communities, where increasing microbial activity and abundance generally correlated with larger grain sizes (Cammen, 1982; Zhang et al., 1998; Cahoon et al., 1999; Musslewhite et al., 2003). Jackson and Weeks (2008) also reported differences in community structure via 16S phylogenetic analysis that were correlated to grain size; specifically, they found that Verrucomicrobia and Planctomycetes were significantly reduced in grain sizes >1000 μm. Albrechtsen and Winding (1992) noted that microbial activity correlated with grain sizes in glacial-fluvio sediments and suggested that this relationship had more to do with sediment permeability than with the grains themselves. More recently, Hemkemeyer et al. (2014) determined through laboratory experiments that both particle size fraction and mineral composition had selective effects on community structure and niche separation.

In our study of Pilot Valley, we also find that grain size is related to, and likely influences discrete microbial community structure, and that this influence relates to multiple geochemical factors. *First*, barium, which is known to adsorb to clays and therefore serves as an indicator of clay concentrations, significantly correlated with Group 1 in the CCA (Figure 5a) (Eylem et al., 1990; Gonneea & Paytan, 2006; Rutten & de Lange, 2002). The dominant taxa in Group 1 (*Halobacteria*, *Salinibacter* and Gemmatimonadetes) all correlate with increasing barium concentration (**Figure 5b**). Both *Halobacteria* and *Salinibacter* require high salt concentrations on the order of 15-30% to thrive, and Gemmatimonadetes is considered a desiccation-resistant phylum (Andrei et al., 2012; DeBruyn et al., 2011; Fawaz, 2013; Oren, 1994; Oren, 2013). The presence of these three taxa in Group 1 makes a case for high salinity pore water in the clay-sized sediment layers of the basin due to low porewater permeability in clays (Bartlett et al., 2010).

*Second*, nitrate concentration was identified as a geochemical factor in both the Mantel correlation and the CCA analyses. Nitrate is only known to accumulate in sediments that have minimal water flow and/or exhibit minimal nitrate-related microbial activity (Lybrand et al., 2013). In the case of Pilot Valley, when the geochemistry of the sub-cores is reorganized by group (**Table 3**), we see that nitrate concentrations are highest in Groups 1 and 3, and below the limit of detection in Group 2. This likely indicates that Group 2 has either higher pore water permeability within this sediment type that results in higher groundwater flow, and/or increased rates of microbial nitrogen cycling.

*Third*, the high sulfate and calcium concentrations associated with Group 2 correlate with large coarse grains that are defined as large euhedral evaporite crystals in **Table 2** (Figure 2). These grains may be mostly composed of gypsum and/or other sulfate salts, hence the elevated calcium and sulfate concentrations may be artifacts of sample preparation for IC and ICP-OES analyses, as these minerals would have dissolved to some degree during the sample extraction process. These large coarse grains allow for greater water movement through the groundwater system and, therefore, continual precipitation and re-dissolution of minerals. Hence, another possible indicator of increased groundwater flow may be the abundance of known SRB in Group 2 (20%) layers, compared to the scarcity of known SRB in Groups 1 and 3 (<2%). Previous studies have suggested that inhibition of sulfate reduction in finer sediments may be due to pore space exclusion of SRB stemming from smaller sediment grain sizes (Bartlett et al., 2010). While mechanical restriction may be a factor in a reduced abundance of SRB in the Group 1 (fine clay) and Group 3 (fine silt) grain types, we feel that to be unlikely, given the suggested presence of similarly-sized microbes such as the *Halobacteria*. By contrast, hydrogen sulfide may exert a more important influence, as sulfate reduction is known to be inhibited by high H_2_S concentrations (Icgen & Harrison, 2006; Okabe et al., 1995; Reis et al., 1992). We hypothesize that compact Group 1 and Group 3 pore water spaces restrict H_2_S diffusion to the extent that SRB cannot thrive in those zones (Roychoudhury et al., 2003), whereas the higher permeability of coarse grains in Group 2 keeps H_2_S from building up to inhibitory concentrations.

### Gemmatimonadetes as a possible indicator of water activity

One of the most interesting aspects of the Pilot Valley microbial community is the high abundance of the phylum Gemmatimonadetes. Originally identified as candidate group BD by Hugenholtz et al. (2001), and as candidate group KS-B by Madrid et al. (Madrid et al., 2001), Gemmatimonadetes was given phylum-level status by Zhang et al. (2003) after that group isolated the first cultured representative, *Gemmatimonas aurantiaca*. On average, members of this phylum represent comprise 2% of soil microbial communities; indeed, Gemmatimonadetes is considered one of the top 9 phyla found in soils (DeBruyn et al., 2011). Members of this enigmatic group are known to possess novel carotenoids as well as photosynthetic capabilities, suggesting an unusual evolutionary history (Takaichi et al., 2010; Zeng et al., 2014). A key characteristic of this phylum is that its members appear to be exceptionally desiccation resistant. Both Debruyn (2011) and Fawaz (2013) provide compelling evidence for their tolerance to low moisture conditions (as low as 8.7%) through biogeographical analyses and laboratory studies across multiple environments. However, the abundance of Gemmatimonadetes in Pilot Valley did not appear to be directly linked with water content, as the difference in water content is small between highest-abundance Group 1(44% Gemmatimonadetes; 24% water content) and lowest-abundance Group 3 (2% Gemmatimonadetes; 28% water content).

Gemmatimonadetes in Pilot Valley co-located with extreme halophilic taxa such as *Halobateria* and *Salinibacter*. This association is intriguing, as no member of the Gemmatimonadetes has yet been shown to be halophilic or halotolerant. Our study provides the first evidence that organisms in the phylum Gemmatimonadetes may not simply be tolerant to desiccation stress (xerotolerance), but also may be tolerant to salinity stress (osmotolerance). A key difference between these two parameters is this: under desiccation stress cellular ionic concentrations increase but generally keep to the same ratios, whereas under salinity stress, the ratios could radically change (Holzinger & Karsten, 2013; Stevenson et al., 2014). Up to the point of this study. As a phylum, Gemmatimonadetes are widely considered to be desiccation-resistant or xerophilic. Our data suggest that at least some Gemmatimonadetes may also be tolerant of high salinity, perhaps even halophilic, potentially defining a novel category of extremophile.

## SUMMARY AND CONCLUSION

In this study, we have identified grain size as a factor that may contribute to the structure of microbial communities in paleolake sediments in Pilot Valley, Utah, a Mars analog environment. This finding has implications from the perspectives of both general microbial ecology and astrobiology. First and foremost, we have shown that microbial communities are more diverse in the vertical than in the horizontal dimension. That is, the evaporitic zonation of the closed basin appears to have little influence on community assembly overall. Secondly, previous studies have suggested that microbial activity in clay sediments effectively ceases due to pore space limitations. However, many of these studies were conducted either prior to advances in high-throughput sequencing of environmental samples, assayed microbial activity in a very limited manner, or did not measure bioactivity at all (Albrechtsen & Winding, 1992; Bartlett et al., 2010; Fredrickson et al., 1997; Li et al., 2015; Zhang et al., 1998). In this study we found that, although sulfate-reducing bacteria are likely inhibited in finer grain and clay-sized sediments, a diverse population of such microbes are nevertheless present, as are a collection of taxa that collectively could carry out the nitrogen cycle. Finally, we have expanded understanding of physiological tolerance within the phylum Gemmatimonadetes, showing that such organisms likely thrive in low-moisture, salty environments.

From an astrobiological perspective, active evaporite mineral formation within a sedimentary system is considered an ideal environment for the preservation and subsequent detection of biosignatures (Summons et al., 2011; Sahl et al., 2008). In Pilot Valley, a system exists that may be actively entraining microbes and, as such, provides a model for learning how to analyze such types of samples. Secondly, the grain size-effects we see on community structure could impact what gets preserved in which sedimentary layers. Preservation would likely be higher in the clay-sized sedimentary layers than in the secondary mineral layers as continual groundwater flow could potentially cause re-dissolution of minerals, destroying biomarkers in those layers. Thirdly, water and the overall moisture content of a system likely selects for organisms specifically capable of metabolism under low water activity, including members of the phylum Gemmatimonadetes.

In closing, Pilot Valley is a fascinating natural laboratory for studying the microbial ecology of hypersaline sediments. Though this study has opened a window onto the phylogenetic diversity of hypersaline sediments, more detailed knowledge of the metabolic diversity of such environments would increase our understanding of how these communities are structured so discretely. Future work will include metagenomic analyses of microbial communities within the Pilot Valley system as well as testing our hypothesis that Pilot Valley Gemmatimonadetes are adapted to thrive in regions of low water activity.

## ACKNOWLEDGEMENTS

This research was supported by funding from the NASA Harriet Jenkins Pre-Doctoral Fellowship Program, the Edna Bailey Sussman Internship Program, the Bechtel K-5 Excellence in Education Initiative, the NASA Astrobiology Institute NNA17BB05A (CAN-7), NASA Astrobiology Institute Director’s Discretionary Fund, The Ford Foundation Fellowship Program, and National Science Foundation Grant NSF IOS 1318843. All DNA sequence data related to this study can be obtained through the European Nucleotide Archive (ENA) via accession number PRJEB11779. The authors thank Drs. Chase Williamson and Lisa Gallagher for training and support on 454 sequencing preparation. The authors would also like to thank Dean Heil and Dr. Jim Ranville for providing access to their respective IC and ICP-OES instruments at Colorado School of Mines and for support of Pilot Valley samples analyzed on those instruments. Finally, the authors thank Drs. Jackson Z. Lee, William Orsi and Joseph Russell III for thoughtful discussions on bioinformatics, multivariate statistics, and coding.

## References

Albrechtsen H-J, Winding A (1992) Microbial biomass and activity in subsurface sediments from Vejen, Denmark. Microb Ecol, 23, 303–317.

Andrei A-Ş, Banciu HL, Oren A (2012) Living with salt: metabolic and phylogenetic diversity of archaea inhabiting saline ecosystems. FEMS Microbiology Letters, 330, 1–9.

Barbieri R, Stivaletta N (2012) Halophiles, Continental Evaporites and the Search for Biosignatures in Environmental Analogues for Mars. Life on Earth and other Planetary Bodies, 24, 13–26.

Barbieri R, Stivaletta N, Marinangeli L, Ori GG (2006) Microbial signatures in sabkha evaporite deposits of Chott el Gharsa (Tunisia) and their astrobiological implications. Planetary and Space Science, 54, 726–736.

Bartlett R, Bottrell SH, Sinclair K, Thornton S, Fielding ID, Hatfield D (2010) Lithological controls on biological activity and groundwater chemistry in Quaternary sediments. Hydrological Processes, 24, 726–735.

Baxter BK, Litchfield CD, Sowers K, Griffith JD, Dassarma PA, Dassarma S (2005) Microbial Diversity of Great Salt Lake. Springer Netherlands, Dordrecht, pp. 9–25.

Boogaerts GL (2015) Preliminary Characterization of the Microbial Community in the Bonneville Salt Flats. In: Department of Biology. University of Alabama at Birmingham, Birmingham, AL.

Boyd ES, Yu R-Q, Barkay T, Hamilton TL, Baxter BK, Naftz DL, Marvin-Dipasquale M (2017) Effect of salinity on mercury methylating benthic microbes and their activities in Great Salt Lake, Utah. Science of The Total Environment, 581-582, 495–506.

Cahoon LB, Nearhoof JE, Tilton CL (1999) Sediment grain size effect on benthic microalgal biomass in shallow aquatic ecosystems. Estuaries, 22, 735–741.

Cammen LM (1982) Effect of Particle Size on Organic Content and Microbial Abundance within Four Marine Sediments. Marine Ecology - Progress Series, 9, 273–280.

Caporaso JG, Bittinger K, Bushman FD, Desantis TZ, Andersen GL, Knight R (2010) PyNAST: a flexible tool for aligning sequences to a template alignment. Bioinformatics, 26, 266–267.

Carling GT, Mayo AL, Tingey D, Bruthans J (2012) Mechanisms, timing, and rates of arid region mountain front recharge. Journal of Hydrology, 428,Äì429, 15–31.

Choi A, Song J, Joung Y, Kogure K, Cho J-C (2015) Lentisphaera profundi sp. nov., isolated from deep-sea water. International Journal of Systematic and Evolutionary Microbiology, 65, 4186–4190.

Couradeau E, Benzerara K, Moreira D, Gérard E, Kaźmierczak J, Tavera R, López-García P (2011) Prokaryotic and Eukaryotic Community Structure in Field and Cultured Microbialites from the Alkaline Lake Alchichica (Mexico). PLoS ONE, 6, e28767.

Currey DR (1990) Quaternary palaeolakes in the evolution of semidesert basins, with special emphasis on Lake Bonneville and the Great Basin, U.S.A. Palaeogeography, Palaeoclimatology, Palaeoecology, 76, 189–214.

Debruyn JM, Nixon LT, Fawaz MN, Johnson AM, Radosevich M (2011) Global Biogeography and Quantitative Seasonal Dynamics of Gemmatimonadetes in Soil. Applied and Environmental Microbiology, 77, 6295–6300.

Delmont TO, Quince C, Shaiber A, Esen ÖC, Lee STM, Rappé MS, Mclellan SL, Lücker S, Eren AM (2018) Nitrogen-fixing populations of Planctomycetes and Proteobacteria are abundant in surface ocean metagenomes. Nature Microbiology, 3, 804–813.

Derito CM, Madsen EL (2008) Stable isotope probing reveals Trichosporon yeast to be active in situ in soil phenol metabolism. ISME Journal, 3, 477–485.

Douglas S (2004) Microbial biosignatures in evaporite deposits: Evidence from Death Valley, California. Planetary and Space Science, 52, 223–227.

Douglas S, Yang H (2002) Mineral biosignatures in evaporites: Presence of rosickyite in an endoevaporitic microbial community from Death Valley, California. Geology, 30, 1075–1078.

Eardley AJ, Gvosdetsky V, Marsell RE (1957) Hydrology of Lake Bonneville and Sediments and Soils of its Basin. Geological Society of America Bulletin, 68, 1141–1201.

Eugster HP, Jones BF (1979) Behavior of major solutes during closed-basin brine evolution. American Journal of Science, 279, 609–631.

Eylem C, Erten HN, Göktürk H (1990) Sorption-desorption behaviour of barium on clays. Journal of Environmental Radioactivity, 11, 183–200.

Fawaz MN (2013) Revealing the Ecological Role of Gemmatimonadetes Through Cultivation and Molecular Analysis of Agricultural Soils. University of Tennessee, Knoxville, TN.

Feazel LM, Spear JR, Berger AB, Harris JK, Frank DN, Ley RE, Pace NR (2008) Eucaryotic Diversity in a Hypersaline Microbial Mat. Applied and Environmental Microbiology, 74, 329–332.

Fornari M, Risacher F, Féraud G (2001) Dating of paleolakes in the central Altiplano of Bolivia. Palaeogeography, Palaeoclimatology, Palaeoecology, 172, 269–282.

Fredrickson JK, Mckinley JP, Bjornstad BN, Long PE, Ringelberg DB, White DC, Krumholz LR, Suflita JM, Colwell FS, Lehman RM, Phelps TJ, Onstott TC (1997) Pore-size constraints on the activity and survival of subsurface bacteria in a late cretaceous shale-sandstone sequence, northwestern New Mexico. Geomicrobiology Journal, 14, 183–202.

Genderjahn S, Alawi M, Mangelsdorf K, Horn F, Wagner D (2018) Desiccation- and Saline-Tolerant Bacteria and Archaea in Kalahari Pan Sediments. Frontiers in microbiology, 9, 2082–2082.

Gonneea ME, Paytan A (2006) Phase associations of barium in marine sediments. Marine Chemistry, 100, 124–135.

Gtari M, Ghodhbane-Gtari F, Nouioui I, Beauchemin N, Tisa LS (2012) Phylogenetic perspectives of nitrogen-fixing actinobacteria. Archives of Microbiology, 194, 3–11.

Hammer Ø, Harper DaT, Ryan PD (2001) PAST: Paleontological statistics software package for education and data analysis. In: Palaeontologia Electronica pp. 9.

Hemkemeyer M, Martens R, Tebbe CC, Heister K, Pronk GJ, Kögel-Knabner I (2014) Artificial soil studies reveal domain-specific preferences of microorganisms for the colonisation of different soil minerals and particle size fractions. FEMS Microbiology Ecology, 90, 770–782.

Heylen K, Keltjens J (2012) Redundancy and modularity in membrane-associated dissimilatory nitrate reduction in Bacillus. Frontiers in Microbiology, 3.

Hollister EB, Engledow AS, Hammett AJM, Provin TL, Wilkinson HH, Gentry TJ (2010) Shifts in microbial community structure along an ecological gradient of hypersaline soils and sediments. ISME J, 4, 829–838.

Holzinger A, Karsten U (2013) Desiccation stress and tolerance in green algae: consequences for ultrastructure, physiological and molecular mechanisms. Frontiers in plant science, 4, 327–327.

Hugenholtz P, Tyson GW, Webb RI, Wagner AM, Blackall LL (2001) Investigation of Candidate Division TM7, a Recently Recognized Major Lineage of the Domain Bacteria with No Known Pure-Culture Representatives. Applied and Environmental Microbiology, 67, 411–419.

Hunt CB (1975) Death Valley: Geology, ecology, archaeology, University of California Press (Berkeley).

Icgen B, Harrison S (2006) Exposure to sulfide causes populations shifts in sulfate-reducing consortia. Research in Microbiology, 157, 784–791.

Jackson CR, Weeks AQ (2008) Influence of particle size on bacterial community structure in aquatic sediments as revealed by 16S rRNA gene sequence analysis. Applied and environmental microbiology, 74, 5237–5240.

Jones BF, White WW, Conko KM, Webster DM, Kohler JF (2009) Mineralogy and Fluid Chemistry of Surficial Sediements in the Newfoundland Basin, Tooele and Box Elder Counties, Utah. Utah Geological Survey, Salt Lake City.

Kjeldsen KU, Loy A, Jakobsen TF, Thomsen TR, Wagner M, Ingvorsen K (2007) Diversity of sulfate-reducing bacteria from an extreme hypersaline sediment, Great Salt Lake (Utah).

Kloppmann W, Négrel P, Casanova J, Klinge H, Schelkes K, Guerrot C (2001) Halite dissolution derived brines in the vicinity of a Permian salt dome (N German Basin). Evidence from boron, strontium, oxygen, and hydrogen isotopes. Geochimica et Cosmochimica Acta, 65, 4087–4101.

Kottek M, Grieser J, Beck C, Rudolf B, Rubel F (2006) World Map of the Koppen-Geiger climate classification updated. Meteorologische Zeitschrift, 15, 259–263.

Legendre P, Fortin M-J (2010) Comparison of the Mantel test and alternative approaches for detecting complex multivariate relationships in the spatial analysis of genetic data. Molecular Ecology Resources, 10, 831–844.

Ley RE, Harris JK, Wilcox J, Spear JR, Miller SR, Bebout BM, Maresca JA, Bryant DA, Sogin ML, Pace NR (2006) Unexpected Diversity and Complexity of the Guerrero Negro Hypersaline Microbial Mat. Applied and Environmental Microbiology, 72, 3685–3695.

Li N, Feng Z, Huang H, Wang X, Dong Z (2015) Lithological and diagenetic restrictions on biogenic gas generation in Songliao Basin inferred from grain size distribution and permeability measurement. Bulletin of Canadian Petroleum Geology, 63, 66–74.

Lindsay MR, Anderson C, Fox N, Scofield G, Allen J, Anderson E, Bueter L, Poudel S, Sutherland K, Munson-Mcgee JH, Van Nostrand JD, Zhou J, Spear JR, Baxter BK, Lageson DR, Boyd ES (2017) Microbialite response to an anthropogenic salinity gradient in Great Salt Lake, Utah. Geobiology, 15, 131–145.

Lines GC (1979) Hydrology and Surface Morphology of the Bonneville Salt Flats and Pilot Valley, Utah. (ed Usgs). U.S. Government Printing Office Washington D.C.

Lozupone CA, Hamady M, Kelley ST, Knight R (2007) Quantitative and Qualitative β Diversity Measures Lead to Different Insights into Factors That Structure Microbial Communities. Applied and Environmental Microbiology, 73, 1576–1585.

Lybrand RA, Michalski G, Graham RC, Parker DR (2013) The geochemical associations of nitrate and naturally formed perchlorate in the Mojave Desert, California, USA. Geochimica et Cosmochimica Acta, 104, 136–147.

Lynch KL, Horgan BH, Munakata-Marr J, Hanley J, Schneider RJ, Rey KA, Spear JR, Jackson WA, Ritter SM (2015) Near-infrared spectroscopy of lacustrine sediments in the Great Salt Lake Desert: An analog study for Martian paleolake basins. Journal of Geophysical Research: Planets, 120, 599–623.

Lynch KL, Jackson WA, Rey K, Spear JR, Rosenzweig F, Munakata Marr J (2019) Evidence for biotic perchlorate reduction in naturally perchlorate-rich sediments of Pilot Valley Basin, Utah. Astrobiology (In Press).

Madrid VM, Aller JY, Aller RC, Chistoserdov AY (2001) High prokaryote diversity and analysis of community structure in mobile mud deposits off French Guiana: identification of two new bacterial candidate divisions.

Madsen DB, Rhode D, Grayson DK, Broughton JM, Livingston SD, Hunt J, Quade J, Schmitt DN, Shaver Iii MW (2001) Late Quaternary environmental change in the Bonneville basin, western USA. Palaeogeography, Palaeoclimatology, Palaeoecology, 167, 243–271.

Mason JL, Kipp KL (1997) Investigation of Salt Loss from the Bonneville Salt Flats, Northwestern Utah. (ed Usgs).

Mcgonigle JM, Bernau JA, Bowen BB, Brazelton W (2019) Robust archaeal and bacterial communities inhabit shallow subsurface sediments of the Bonneville Salt Flats. bioRxiv, 553032.

Meuser J, Baxter B, Spear J, Peters J, Posewitz M, Boyd E (2013) Contrasting Patterns of Community Assembly in the Stratified Water Column of Great Salt Lake, Utah. Microb Ecol, 66, 268–280.

Mohammadi SS, Pol A, Van Alen T, Jetten MSM, Op Den Camp HJM (2017) Ammonia Oxidation and Nitrite Reduction in the Verrucomicrobial Methanotroph Methylacidiphilum fumariolicum SolV. Frontiers in Microbiology, 8.

Morin N, Vallaeys T, Hendrickx L, Natalie L, Wilmotte A (2010) An efficient DNA isolation protocol for filamentous cyanobacteria of the genus Arthrospira. Journal of Microbiological Methods, 80, 148–154.

Musslewhite CL, Mcinerney MJ, Dong H, Onstott TC, Green-Blum M, Swift D, Macnaughton S, White DC, Murray C, Chien Y-J (2003) The Factors Controlling Microbial Distribution and Activity in the Shallow Subsurface Geomicrobiology Journal, 20, 245–261.

Myers MR, King GM (2017) Perchlorate-Coupled Carbon Monoxide (CO) Oxidation: Evidence for a Plausible Microbe-Mediated Reaction in Martian Brines. Frontiers in Microbiology, 8.

Okabe S, Nielsen PH, Jones WL, Characklis WG (1995) Sulfide product inhibition of Desulfovibrio desulfuricans in batch and continuous cultures. Water Research, 29, 571–578.

Oren A (1994) The ecology of the extremely halophilic archaea. FEMS Microbiology Reviews, 13, 415–439.

Oren A (2006) Life at High Salt Concentrations. In: The Prokaryotes (eds Dworkin M, Falkow S, Rosenberg E, Schleifer K-H, Stackebrandt E). Springer New York, pp. 263–282.

Oren A (2008) Microbial life at high salt concentrations: phylogenetic and metabolic diversity. Saline Systems, 4, 2.

Oren A (2013) Salinibacter: an extremely halophilic bacterium with archaeal properties.

Osburn MR, Sessions AL, Pepe-Ranney C, Spear JR (2011) Hydrogen-isotopic variability in fatty acids from Yellowstone National Park hot spring microbial communities. Geochimica et Cosmochimica Acta, 75, 4830–4845.

Oviatt CG (2015) Chronology of Lake Bonneville, 30,000 to 10,000 yr B.P. Quaternary Science Reviews, 110, 166–171.

Post FJ (1977) The microbial ecology of the Great Salt Lake. Microb Ecol, 3, 143–165.

Reis MaM, Almeida JS, Lemos PC, Carrondo MJT (1992) Effect of hydrogen sulfide on growth of sulfate reducing bacteria. Biotechnology and Bioengineering, 40, 593–600.

Rey KA, Mayo AL, Tingey DG, Nelson ST (2016) Chapter 10 - Late Pleistocene to Early Holocene Sedimentary History of the Lake Bonneville Pilot Valley Embayment, Utah-Nevada, USA. In: Developments in Earth Surface Processes (eds Oviatt CG, Shroder JF). Elsevier, pp. 184–220.

Robertson CE, Spear JR, Harris JK, Pace NR (2009) Diversity and Stratification of Archaea in a Hypersaline Microbial Mat. Applied and Environmental Microbiology, 75, 1801–1810.

Roychoudhury AN, Van Cappellen P, Kostka JE, Viollier E (2003) Kinetics of microbially mediated reactions: dissimilatory sulfate reduction in saltmarsh sediments (Sapelo Island, Georgia, USA). Estuarine, Coastal and Shelf Science, 56, 1001–1010.

Rude PD, Aller RC (1991) Fluorine mobility during early diagenesis of carbonate sediment: An indicator of mineral transformations. Geochimica et Cosmochimica Acta, 55, 2491–2509.

Rutten A, De Lange GJ (2002) A novel selective extraction of barite, and its application to eastern Mediterranean sediments. Earth and Planetary Science Letters, 198, 11–24.

Sahl JW, Pace NR, Spear JR (2008) Comparative Molecular Analysis of Endoevaporitic Microbial Communities. Applied and Environmental Microbiology, 74, 6444–6446.

Schwendner P, Bohmeier M, Rettberg P, Beblo-Vranesevic K, Gaboyer F, Moissl-Eichinger C, Perras AK, Vannier P, Marteinsson VT, Garcia-Descalzo L, Gómez F, Malki M, Amils R, Westall F, Riedo A, Monaghan EP, Ehrenfreund P, Cabezas P, Walter N, Cockell C (2018) Beyond Chloride Brines: Variable Metabolomic Responses in the Anaerobic Organism Yersinia intermedia MASE-LG-1 to NaCl and MgSO4 at Identical Water Activity. Frontiers in Microbiology, 9.

Sirisena KA, Ramirez S, Steele A, Glamoclija M (2018) Microbial Diversity of Hypersaline Sediments from Lake Lucero Playa in White Sands National Monument, New Mexico, USA. Microb Ecol, 76, 404–418.

Spencer RJ, Baedecker MJ, Eugster HP, Forester RM, Goldhaber MB, Jones BF, Kelts K, Mckenzie J, Madsen DB, Rettig SL, Rubin M, Bowser CJ (1984) Great Salt Lake, and precursors, Utah: The last 30,000 years. Contributions to Mineralogy and Petrology, 86, 321–334.

Stevenson A, Cray JA, Williams JP, Santos R, Sahay R, Neuenkirchen N, Mcclure CD, Grant IR, Houghton JDR, Quinn JP, Timson DJ, Patil SV, Singhal RS, Antón J, Dijksterhuis J, Hocking AD, Lievens B, Rangel DEN, Voytek MA, Gunde-Cimerman N, Oren A, Timmis KN, Mcgenity TJ, Hallsworth JE (2014) Is there a common water-activity limit for the three domains of life? The Isme Journal, 9, 1333.

Summons RE, Amend JP, Bish D, Buick R, Cody GD, Des Marais DJ, Dromart G, Eigenbrode JL, Knoll AH, Sumner DY (2011) Preservation of Martian Organic and Environmental Records: Final Report of the Mars Biosignature Working Group. Astrobiology, 11, 157–181.

Takaichi S, Maoka T, Takasaki K, Hanada S (2010) Carotenoids of Gemmatimonas aurantiaca (Gemmatimonadetes): identification of a novel carotenoid, deoxyoscillol 2-rhamnoside, and proposed biosynthetic pathway of oscillol 2,2’-dirhamnoside. Microbiology, 156, 757–763.

Tazi L, Breakwell DP, Harker AR, Crandall KA (2014) Life in extreme environments: microbial diversity in Great Salt Lake, Utah. Extremophiles, 18, 525–535.

Turk LJ, Davis SN, Bingham CP (1973) Hydrogeology of Lacustrine Sediments, Bonneville Salt Flats, Utah. Economic Geology, 68, 65–78.

Ventosa A, Arahal DR (2009) Physico-chemical characteristics of hypersaline environments and their biodiversity. In: Extremophiles (eds Gerday C, Glansdorff N). Eolss Publishers, Paris, France, pp. 242–262.

Ventosa A, Mellado E, Sanchez-Porro C, Marquez MC (2008) Halophilic and Halotolerant Micro-Organisms from Soils. In: Microbiology of Extreme Soils (eds Dion P, Nautiyal CS). Springer, Berlin, pp. 87–115.

Ward NL, Challacombe JF, Janssen PH, Henrissat B, Coutinho PM, Wu M, Xie G, Haft DH, Sait M, Badger J, Barabote RD, Bradley B, Brettin TS, Brinkac LM, Bruce D, Creasy T, Daugherty SC, Davidsen TM, Deboy RT, Detter JC, Dodson RJ, Durkin AS, Ganapathy A, Gwinn-Giglio M, Han CS, Khouri H, Kiss H, Kothari SP, Madupu R, Nelson KE, Nelson WC, Paulsen I, Penn K, Ren Q, Rosovitz MJ, Selengut JD, Shrivastava S, Sullivan SA, Tapia R, Thompson LS, Watkins KL, Yang Q, Yu C, Zafar N, Zhou L, Kuske CR (2009) Three Genomes from the Phylum *Acidobacteria* Provide Insight into the Lifestyles of These Microorganisms in Soils. Applied and Environmental Microbiology, 75, 2046–2056.

Warren JK (1999) Evaporites: Their Evolution and Economics, Blackwell Science, Oxford.

Wilkerson C (2012) Utah’s Great Salt Lake and Ancient Lake Bonneville. In: Lake Bonneville. Utah Geological Survey.

Wrcc (2013) General Climate Summary Records. Wester Regional Climate Center.

Zeng Y, Feng F, Medová H, Dean J, Koblížek M (2014) Functional type 2 photosynthetic reaction centers found in the rare bacterial phylum Gemmatimonadetes. Proceedings of the National Academy of Sciences, 111, 7795–7800.

Zhang C, Palumbo AV, Phelps TJ, Beauchamp JJ, Brockman FJ, Murray CJ, Parsons BS, Swift DJP (1998) Grain size and depth constraints on microbial variability in coastal plain subsurface sediments. Geomicrobiology Journal, 15, 171–185.

Zhang H, Sekiguchi Y, Hanada S, Hugenholtz P, Kim H, Kamagata Y, Nakamura K (2003) Gemmatimonas aurantiaca gen. nov., sp. nov., a Gram-negative, aerobic, polyphosphate-accumulating micro-organism, the first cultured representative of the new bacterial phylum Gemmatimonadetes phyl. nov. International Journal of Systematic and Evolutionary Microbiology, 53, 1155–1163.

