## Supplemental File for "Discrete Community Assemblages Within Hypersaline Paleolake Sediments of Pilot Valley, Utah"

**for**

1. **SI Tables**

**Table S1.** Summary 454 Sequencing Statistics

**Table S2.** Detailed Pilot Valley Geochemistry - Sediment Cations

**Table S3.** Detailed Pilot Valley Geochemistry - Sediment Anions

**Table S4.** Detailed Pilot Valley Geochemistry - Hydration and Carbon

**Table S5.** Pilot Valley Aquifer Brine Chemistry

**Table S6.** Summary Anions for Comparison Study

**Table S7.** Mantel Correlations for Transect Study

**Table S8.** Detailed Pilot Valley Geochemistry – Sediment Cations by Group

**Table S9**. Detailed Pilot Valley Geochemistry – Sediment Anions by Group

**Table S10**. Detailed Pilot Valley Geochemistry – Hydration and Carbon by Group

2. **SI Figures**

**Figure S1.** Rarefaction plots of for Field Site Comparison Study

**Figure S2.** PCoA plots for Comparison Study of a)Bray-Curtis, b)un-weighted	Unifrac, and c) weighted Unifrac beta diversity metrics.

**Figure S3.** Rarefaction plots for Pilot Valley Transect Study

**Supplementary Data Tables**

| Table S1. Summary Sequencing Statistics | | | | | | | | | |
| --- | --- | --- | --- | --- | --- | --- | --- | --- | --- |
| Study | Total Raw Sequences* | Total QC Sequences | Mean Sequence Length | Sequence Number | | | | |  |
|  |  |  |  | Min | Max | Median | Mean | SD | #OTU's |
| Comparison | 89,827 | 55,754 | 394 | 8028 | 14,223 | 11,771 | 11,151 | 2672 | 1769 |
| Transect | 209,045 | 190,806 | 382 | 6580 | 16,178 | 7634 | 8296 | 2012 | 1336 |

*Raw sequences after "split_library.py" command script to parse sequences from shared 454 plates.

| Table S2. Detailed Pilot Valley Geochemistry - Sediment Cations  BDL=Below Detection Limit ND=No Data | | | | | | | | | | |
| --- | --- | --- | --- | --- | --- | --- | --- | --- | --- | --- |
| Core ID | Ca | Na | K | Mg | Fe | Ba | Li | Sr | V | Zn |
|  | wt % | wt % | wt% | wt% | mg/kg | mg/kg | mg/kg | mg/kg | mg/kg | mg/kg |
| **PV 1** |  |  |  |  |  |  |  |  |  |  |
| PV1-1-3 | 0.04 | 2.64 | 0.16 | 0.05 | BDL | 9.99 | 22.3 | 71.4 | BDL | 6.58 |
| PV1-4 | 0.04 | 2.42 | 0.15 | 0.04 | BDL | 6.08 | 22.8 | 14.5 | BDL | BDL |
| PV1-5 | 0.03 | 2.36 | 0.15 | 0.03 | BDL | 3.46 | 19.4 | 9.5 | BDL | BDL |
| ***Average*** | ***0.04*** | ***2.47*** | ***0.15*** | ***0.04*** | ***BDL*** | ***6.51*** | ***21.5*** | ***31.8*** | ***BDL*** | ***6.58*** |
| **PV 2** |  |  |  |  |  |  |  |  |  |  |
| PV2-1 | 1.55 | 2.22 | 0.16 | 0.07 | 1.92 | 0.17 | 20.9 | 123 | 0.89 | 3.97 |
| PV2-2 | 0.05 | 2.44 | 0.15 | 0.04 | BDL | 3.35 | 20.3 | 36.2 | 0.53 | BDL |
| PV2-3 | 0.20 | 2.94 | 0.21 | 0.07 | 2.78 | 6.15 | 29.1 | 61.3 | 1.21 | 3.22 |
| PV2-4 | 0.08 | 2.91 | 0.18 | 0.06 | BDL | 5.31 | 26.6 | 48.1 | 1.50 | BDL |
| PV2-5 | 1.23 | 2.72 | 0.20 | 0.08 | BDL | 5.12 | 24.6 | 103 | 0.63 | 2.99 |
| PV2-6 | 1.31 | 2.42 | 0.18 | 0.07 | BDL | 2.94 | 23.9 | 51.7 | 0.67 | 3.89 |
| PV2-7 | 1.41 | 1.60 | 0.10 | 0.05 | BDL | 3.55 | 16.5 | 233 | BDL | 3.40 |
| PV2-8 | 1.03 | 3.02 | 0.20 | 0.10 | BDL | 5.22 | 29.7 | 356 | BDL | 2.83 |
| PV2-9 | 1.51 | 2.45 | 0.16 | 0.08 | BDL | 2.81 | 22.7 | 191 | BDL | 3.93 |
| PV2-10 | 0.01 | 3.27 | 0.20 | 0.09 | 1.35 | 0.28 | 27.4 | 40.4 | 0.96 | 2.25 |
| PV2-11 | 0.09 | 4.64 | 0.30 | 0.12 | 1.31 | 1.01 | 49.1 | 21.6 | 1.88 | 4.54 |
| ***Average*** | ***0.77*** | ***2.78*** | ***0.19*** | ***0.08*** | ***1.84*** | ***3.26*** | ***26.4*** | ***115*** | ***1.03*** | ***3.45*** |
| **PV 3** |  |  |  |  |  |  |  |  |  |  |
| PV3-1 | 1.40 | 1.89 | 0.35 | 0.09 | BDL | 1.90 | 26.5 | 90.3 | BDL | 3.84 |
| PV3-2 | 1.25 | 1.94 | 0.41 | 0.09 | BDL | 3.28 | 27.3 | 79.1 | BDL | 3.82 |
| PV3-3 | 1.24 | 0.70 | 0.09 | 0.03 | BDL | 1.99 | 7.82 | 32.8 | BDL | 3.38 |
| ***Average*** | ***1.30*** | ***1.51*** | ***0.28*** | ***0.07*** | ***BDL*** | ***2.39*** | ***20.5*** | ***67.4*** | ***BDL*** | ***3.68*** |
| **PV 5** |  |  |  |  |  |  |  |  |  |  |
| PV5-1 | ND | ND | ND | ND | ND | ND | ND | ND | ND | ND |
| PV5-2 | 1.29 | 1.15 | 0.15 | 0.04 | BDL | 1.43 | 10.0 | 64.9 | BDL | 3.65 |
| PV5-3 | 1.29 | 1.32 | 0.14 | 0.03 | BDL | 1.58 | 8.87 | 80.9 | BDL | 3.77 |
| PV5-4 | 1.40 | 2.22 | 0.38 | 0.07 | BDL | 4.01 | 20.8 | 213 | BDL | 4.06 |
| ***Average*** | ***1.33*** | ***1.56*** | ***0.22*** | ***0.05*** | ***BDL*** | ***2.34*** | ***13.2*** | ***119*** | ***BDL*** | ***3.82*** |

| Table S3. Detailed Pilot Valley Geochemistry - Sediment Anions  BDL=Below Detection Limit ND=No Data | | | | | | | |
| --- | --- | --- | --- | --- | --- | --- | --- |
| Core ID | SO_4_- | Cl | NO_3_- | Br | F | ClO_3_- | ClO_4_- |
|  | wt % | wt % | mg/kg | mg/kg | mg/kg | μg/kg | μg/kg |
| **PV 1** |  |  |  |  |  |  |  |
| PV1-1-3 | 0.11 | 3.99 | 16.1 | BDL | BDL | 1.85 | 0.82 |
| PV1-4 | 0.08 | 3.77 | 14.4 | BDL | BDL | 0.85 | 0.64 |
| PV1-5 | 0.14 | 3.66 | 10.0 | BDL | 7.90 | ND | ND |
| ***Average*** | ***0.11*** | ***3.81*** | ***13.50*** | ***BDL*** | ***7.90*** | ***1.35*** | ***0.73*** |
| **PV 2** |  |  |  |  |  |  |  |
| PV2-1 | 3.35 | 3.08 | 3.70 | BDL | BDL | 0.44 | 0.47 |
| PV2-2 | 0.11 | 4.14 | 4.30 | BDL | BDL | 0.99 | 0.92 |
| PV2-3 | 0.52 | 4.68 | 4.50 | BDL | BDL | 0.40 | 0.56 |
| PV2-4 | 0.14 | 4.69 | 3.00 | BDL | BDL | ND | ND |
| PV2-5 | 2.90 | 4.27 | BDL | BDL | BDL | ND | ND |
| PV2-6 | 3.23 | 3.74 | BDL | BDL | 8.10 | ND | ND |
| PV2-7 | 3.29 | 2.46 | BDL | BDL | BDL | ND | ND |
| PV2-8 | 2.43 | 4.87 | BDL | BDL | 9.90 | ND | ND |
| PV2-9 | 3.36 | 3.63 | BDL | BDL | 7.60 | ND | ND |
| PV2-10 | 1.67 | 5.03 | BDL | BDL | 8.40 | ND | ND |
| PV2-11 | 0.19 | 7.49 | BDL | BDL | 8.60 | ND | ND |
| ***Average*** | ***1.93*** | ***4.37*** | ***3.88*** | ***BDL*** | ***8.52*** | ***0.61*** | ***0.65*** |
| **PV 3** |  |  |  |  |  |  |  |
| PV3-1 | 3.80 | 4.21 | 19.8 | 24.1 | 14.2 | 0.69 | 0.44 |
| PV3-2 | 3.38 | 4.15 | BDL | 22.8 | 1.60 | 0.69 | 0.44 |
| PV3-3 | 3.21 | 1.58 | BDL | 8.60 | BDL | 0.57 | 0.30 |
| ***Average*** | ***3.46*** | ***3.31*** | ***19.80*** | ***18.50*** | ***7.90*** | ***0.65*** | ***0.39*** |
| **PV 5** |  |  |  |  |  |  |  |
| PV5-1 | ND | ND | ND | ND | ND | ND | ND |
| PV5-2 | 3.32 | 2.49 | BDL | 15.3 | BDL | 1.34 | 0.34 |
| PV5-3 | 3.24 | 2.86 | 2.10 | 14.5 | 2.50 | 0.42 | 0.24 |
| PV5-4 | 3.85 | 4.69 | BDL | 15.8 | 1.70 | 0.30 | 0.26 |
| ***Average*** | ***3.47*** | ***3.35*** | ***2.10*** | ***15.20*** | ***2.10*** | ***0.68*** | ***0.28*** |
| PV5-1 | ND | ND | ND | ND | ND | ND | ND |
| PV5-2 | 3.32 | 2.49 | BDL | 15.3 | BDL | 1.34 | 0.34 |
| PV5-3 | 3.24 | 2.86 | 2.10 | 14.5 | 2.50 | 0.42 | 0.24 |
| PV5-4 | 3.85 | 4.69 | BD | 15.8 | 1.70 | 0.30 | 0.26 |
| ***Average*** | ***3.47*** | ***3.35*** | ***2.10*** | ***15.20*** | ***2.10*** | ***0.68*** | ***0.28*** |

| Table S4. Detailed Pilot Valley Geochemistry - Hydration and Carbon  BDL=Below Detection Limit ND=No Data | | | | |
| --- | --- | --- | --- | --- |
| Core ID | TOC | TIC | Total Carbon | Avg. Hydration |
|  | wt% | wt% | wt% | wt% |
| **PV 1** |  |  |  |  |
| PV1-1-3 | 0.61 | 2.36 | 2.97 | 24.10 |
| PV1-4 | 0.28 | 3.25 | 3.52 | 26.60 |
| PV1-5 | 0.42 | 3.85 | 4.27 | 31.60 |
| ***Average*** | ***0.44*** | ***3.15*** | ***3.59*** | ***27.43*** |
| **PV 2** |  |  |  |  |
| PV2-1 | 0.43 | 1.70 | 2.13 | 31.60 |
| PV2-2 | 0.54 | 2.63 | 3.17 | 21.80 |
| PV2-3 | 1.10 | 1.98 | 3.08 | 25.40 |
| PV2-4 | 0.72 | 2.40 | 3.12 | 26.20 |
| PV2-5 | 0.36 | 1.58 | 1.94 | 26.60 |
| PV2-6 | 0.88 | 1.94 | 2.82 | 27.70 |
| PV2-7 | 0.42 | 1.01 | 1.43 | 29.20 |
| PV2-8 | 1.40 | 5.01 | 6.41 | 25.10 |
| PV2-9 | 0.26 | 4.07 | 4.32 | 28.10 |
| PV2-10 | 0.70 | 5.79 | 6.49 | 31.00 |
| PV2-11 | 0.96 | 5.07 | 6.03 | 38.50 |
| ***Average*** | ***0.71*** | ***3.02*** | ***3.72*** | ***28.29*** |
| **PV 3** |  |  |  |  |
| PV3-1 | 0.88 | 0.97 | 1.9 | 27.0 |
| PV3-2 | 0.78 | 1.45 | 2.24 | 23.7 |
| PV3-3 | 0.16 | 0.34 | 0.50 | 23.80 |
| ***Average*** | ***0.61*** | ***0.92*** | ***1.53*** | ***24.83*** |
| **PV 5** |  |  |  |  |
| PV5-1 | ND | ND | ND | ND |
| PV5-2 | 0.13 | 0.28 | 0.40 | 24.1 |
| PV5-3 | 0.24 | 0.01 | 0.25 | 24.2 |
| PV5-4 | 0.75 | 1.67 | 2.42 | 27.0 |
| ***Average*** | ***0.37*** | ***0.65*** | ***1.02*** | ***25.10*** |

| Table S5. Anion Data for Comparison Sites-^a^  BDL=Below Detection Limit ND=No Data | | | | |
| --- | --- | --- | --- | --- |
| Field Site |  | Cl | SO_4_ | NO_3_ |
|  |  | wt% | wt% | mg/kg |
| GSL- SJ |  | 4.26 | 1.15 | BDL |
| BSF-Rim |  | 5.23 | 4.16 | BDL |
| PV-Rim |  | 3.09 | 0.77 | 35.5 |
| ^a^No cation data are available for these samples. BDL = Below Detection Limit | | | | |

| Table S6. Pilot Valley Aquifer Brine Chemistry^a^  BDL=Below Detection Limit ND=No Data | | | | | | | | | |
| --- | --- | --- | --- | --- | --- | --- | --- | --- | --- |
|  | *Cations* | | | |  |  | *Anions* | | |
| Sample Site | Ca | K | Mg | Na | Fe |  | Cl | SO_4_ | NO_3_ |
|  | w/v% | w/v% | w/v% | w/v% | mg/L |  | w/v% | w/v% | mg/L |
| PV-1 | ND | ND | ND | ND | ND |  | ND | ND | ND |
| PV-2 | 0.21 | 0.98 | 0.25 | 8.57 | 0.13 |  | 17.8 | 0.42 | 50.7 |
| PV-3 | 0.26 | 1.40 | 0.31 | 8.83 | BDL |  | 19.2 | 0.27 | 67.2 |
| PV-5 | 0.25 | 1.20 | 0.22 | 9.62 | 1.72 |  | 20.2 | 0.27 | 48.2 |
| ^a^There is no data for Sample Site PV-1 as the bore hole could not reach the water table at the rim of the basin. | | | | | | | | | |

| Table S7. Mantel Correlations | | | | | | | | | |
| --- | --- | --- | --- | --- | --- | --- | --- | --- | --- |
|  |  |  | Weighted Unifrac | | |  | Unweighted Unifrac | | |
| Environmental Variable | |  | (r) |  | *P*-value |  | (r) |  | *P*-value |
| **Ba** | |  | 0.175 |  | 0.015* |  | 0.235 |  | 0.009** |
| Br | |  | -0.040 |  | 0.458 |  | 0.052 |  | 0.422 |
| Ca | |  | 0.047 |  | 0.422 |  | 0.204 |  | 0.013* |
| Cl | |  | -0.254 |  | 0.655 |  | 0.019 |  | 0.793 |
| F | |  | 0.029 |  | 0.601 |  | 0.088 |  | 0.126 |
| Fe | |  | 0.025 |  | 0.640 |  | 0.022 |  | 0.784 |
| K | |  | -0.021 |  | 0.713 |  | -0.042 |  | 0.520 |
| Li | |  | -0.065 |  | 0.253 |  | 0.011 |  | 0.893 |
| Mg | |  | -0.044 |  | 0.455 |  | 0.035 |  | 0.621 |
| Na | |  | -0.022 |  | 0.740 |  | 0.042 |  | 0.570 |
| NO_3-_ | |  | 0.015 |  | 0.802 |  | 0.162 |  | 0.016* |
| SO_4-_ | |  | 0.056 |  | 0.260 |  | 0.208 |  | 0.017* |
| Sr | |  | 0.052 |  | 0.316 |  | 0.059 |  | 0.382 |
| V | |  | -0.017 |  | 0.76 |  | -0.042 |  | 0.507 |
| Zn | |  | -0.026 |  | 0.658 |  | 0.016 |  | 0.81 |
| Water (%) | |  | 0.005 |  | 0.922 |  | 0.091 |  | 0.15 |
| Total Organic Carbon | |  | 0.004 |  | 0.957 |  | -0.055 |  | 0.401 |
| Total Inorganic Carbon | |  | -0.054 |  | 0.410 |  | 0.089 |  | 0.157 |
| Mantel test statistic and P-value based on 999 permutations for each variable. **P*≤0.05, ***P*≤0.01 | | | | | | | | | |

| Table S8. Pilot Valley Geochemistry - Sediment Cations By Group  BDL=Below Detection Limit ND=No Data | | | | | | | | | | | | | | | | | | |  |
| --- | --- | --- | --- | --- | --- | --- | --- | --- | --- | --- | --- | --- | --- | --- | --- | --- | --- | --- | --- |
| Core ID | | Ca | | Na | | K | | Mg | Fe | | Ba | | Li | | Sr | | V | Zn | Mo |
|  |  | wt % | | wt % | | wt% | | wt% | mg/kg | | mg/kg | | mg/kg | | mg/kg | | mg/kg | mg/kg | mg/kg |
| **Group 1** | | | | | | | | | | | | | | | | | | | |
| PV2-2 | | 0.05 | | 2.44 | | 0.15 | | 0.04 | BDL | | 3.35 | | 20.3 | | 36.2 | | 0.53 | BDL | BDL |
| PV2-3 | | 0.20 | | 2.94 | | 0.21 | | 0.07 | 2.78 | | 6.15 | | 29.1 | | 61.3 | | 1.21 | 3.22 | BDL |
| PV2-4 | | 0.08 | | 2.91 | | 0.18 | | 0.06 | BDL | | 5.31 | | 26.6 | | 48.1 | | 1.50 | BDL | BDL |
| PV2-5 | | 1.23 | | 2.72 | | 0.20 | | 0.08 | BDL | | 5.12 | | 24.6 | | 103 | | 0.63 | 2.99 | BDL |
| PV5-4 | | 1.40 | | 2.22 | | 0.38 | | 0.07 | BDL | | 4.01 | | 20.8 | | 213 | | BDL | 4.06 | BDL |
| ***Average*** | | ***0.59*** | | ***2.65*** | | ***0.23*** | | ***0.07*** | ***2.78*** | | ***4.79*** | | ***24.27*** | | ***92.28*** | | ***0.97*** | ***3.42*** | ***BDL*** |
| **Group 2** | | | | | | | | | | | | | | | | | | | |
| PV2-6 | | 1.31 | | 2.42 | | 0.18 | | 0.07 | BDL | | 2.94 | | 23.9 | | 51.7 | | 0.67 | 3.89 | BDL |
| PV2-7 | | 1.41 | | 1.60 | | 0.10 | | 0.05 | BDL | | 3.55 | | 16.5 | | 233 | | BDL | 3.40 | BDL |
| PV2-8 | | 1.03 | | 3.02 | | 0.20 | | 0.10 | BDL | | 5.22 | | 29.7 | | 356 | | BDL | 2.83 | 1.903 |
| PV2-9 | | 1.51 | | 2.45 | | 0.16 | | 0.08 | BDL | | 2.81 | | 22.7 | | 191 | | BDL | 3.93 | BDL |
| PV3-2 | | 1.25 | | 1.94 | | 0.41 | | 0.09 | BDL | | 3.28 | | 27.3 | | 79.1 | | BDL | 3.82 | BDL |
| PV3-3 | | 1.24 | | 0.70 | | 0.09 | | 0.03 | BDL | | 1.99 | | 7.82 | | 32.8 | | BDL | 3.38 | BDL |
| ***Average*** | | ***1.29*** | | ***2.02*** | | ***0.19*** | | ***0.07*** | ***BDL*** | | ***3.30*** | | ***21.31*** | | ***157.37*** | | ***0.67*** | ***3.54*** | ***1.90*** |
| **Group 3** | | | | | | | | | | | | | | | | | | | |
| PV1-4 | | 0.04 | | 2.42 | | 0.15 | | 0.04 | BDL | | 6.08 | | 22.8 | | 14.5 | | BDL | BDL | BDL |
| PV1-5 | | 0.03 | | 2.36 | | 0.15 | | 0.03 | BDL | | 3.46 | | 19.4 | | 9.5 | | BDL | BDL | BDL |
| PV2-1 | | 1.55 | | 2.22 | | 0.16 | | 0.07 | 1.92 | | 0.17 | | 20.9 | | 123 | | 0.89 | 3.97 | BDL |
| PV2-10 | | 0.01 | | 3.27 | | 0.20 | | 0.09 | 1.35 | | 0.28 | | 27.4 | | 40.4 | | 0.96 | 2.25 | BDL |
| PV2-11 | | 0.09 | | 4.64 | | 0.30 | | 0.12 | 1.31 | | 1.01 | | 49.1 | | 21.6 | | 1.88 | 4.54 | 8.299 |
| PV3-1 | | 1.40 | | 1.89 | | 0.35 | | 0.09 | BDL | | 1.90 | | 26.5 | | 90.3 | | BDL | 3.84 | BDL |
| PV5-1 | | ND | | ND | | ND | | ND | ND | | ND | | ND | | ND | | ND | ND | ND |
| PV5-2 | | 1.29 | | 1.15 | | 0.15 | | 0.04 | BDL | | 1.43 | | 10.0 | | 64.9 | | BDL | 3.65 | BDL |
| PV5-3 | | 1.29 | | 1.32 | | 0.14 | | 0.03 | BDL | | 1.58 | | 8.87 | | 80.9 | | BDL | 3.77 | BDL |
| ***Average*** | | ***0.71*** | | ***2.41*** | | ***0.20*** | | ***0.07*** | ***1.53*** | | ***1.99*** | | ***23.12*** | | ***55.69*** | | ***1.25*** | ***3.67*** | ***8.30*** |
| Table S9. Detailed Pilot Valley Geochemistry - Sediment Anions By Group  BDL=Below Detection Limit ND=No Data | | | | | | | | | | | | | | | |  |  |  |  |
| Core ID | SO_4_- | | Cl | | NO_3_- | | Br | | | F | | ClO_3_- | | ClO_4_- | |  |  |  |  |
|  | wt % | | wt % | | mg/kg | | mg/kg | | | mg/kg | | μg/kg | | μg/kg | |  |  |  |  |
| **Group 1** | | | | | | | | | | | | | | | |  |  |  |  |
| PV2-2 | 0.11 | | 4.14 | | 4.30 | | BDL | | | BDL | | 0.99 | | 0.92 | |  |  |  |  |
| PV2-3 | 0.52 | | 4.68 | | 4.50 | | BDL | | | BDL | | 0.40 | | 0.56 | |  |  |  |  |
| PV2-4 | 0.14 | | 4.69 | | 3.00 | | BDL | | | BDL | | ND | | ND | |  |  |  |  |
| PV2-5 | 2.90 | | 4.27 | | BDL | | BDL | | | BDL | | ND | | ND | |  |  |  |  |
| PV5-4 | 3.85 | | 4.69 | | BDL | | 15.8 | | | 1.70 | | 0.30 | | 0.26 | |  |  |  |  |
| ***Average*** | ***1.50*** | | ***4.49*** | | ***3.93*** | | ***15.80*** | | | ***1.70*** | | ***0.56*** | | ***0.58*** | |  |  |  |  |
| **Group 2** | | | | | | | | | | | | | | | |  |  |  |  |
| PV2-6 | 3.23 | | 3.74 | | BDL | | BDL | | | 8.10 | | ND | | ND | |  |  |  |  |
| PV2-7 | 3.29 | | 2.46 | | BDL | | BDL | | | BDL | | ND | | ND | |  |  |  |  |
| PV2-8 | 2.43 | | 4.87 | | BDL | | BDL | | | 9.90 | | ND | | ND | |  |  |  |  |
| PV2-9 | 3.36 | | 3.63 | | BDL | | BDL | | | 7.60 | | ND | | ND | |  |  |  |  |
| PV3-2 | 3.38 | | 4.15 | | BDL | | 22.8 | | | 1.60 | | 0.69 | | 0.44 | |  |  |  |  |
| PV3-3 | 3.21 | | 1.58 | | BDL | | 8.60 | | | BDL | | 0.57 | | 0.30 | |  |  |  |  |
| ***Average*** | ***3.15*** | | ***3.41*** | | ***BDL*** | | ***15.70*** | | | ***6.80*** | | ***0.63*** | | ***0.37*** | |  |  |  |  |
| **Group 3** | | | | | | | | | | | | | | | |  |  |  |  |
| PV1-4 | 0.08 | | 3.77 | | 14.4 | | BDL | | | BDL | | 0.85 | | 0.64 | |  |  |  |  |
| PV1-5 | 0.14 | | 3.66 | | 10.0 | | BDL | | | 7.90 | | ND | | ND | |  |  |  |  |
| PV2-1 | 3.35 | | 3.08 | | 3.70 | | BDL | | | BDL | | 0.44 | | 0.47 | |  |  |  |  |
| PV2-10 | 1.67 | | 5.03 | | BDL | | BDL | | | 8.40 | | ND | | ND | |  |  |  |  |
| PV2-11 | 0.19 | | 7.49 | | BDL | | BDL | | | 8.60 | | ND | | ND | |  |  |  |  |
| PV3-1 | 3.80 | | 4.21 | | 19.8 | | 24.1 | | | 14.2 | | 0.69 | | 0.44 | |  |  |  |  |
| PV5-2 | 3.32 | | 2.49 | | BDL | | 15.3 | | | BDL | | 1.34 | | 0.34 | |  |  |  |  |
| PV5-3 | 3.24 | | 2.86 | | 2.10 | | 14.5 | | | 2.50 | | 0.42 | | 0.24 | |  |  |  |  |
| ***Average*** | ***1.97*** | | ***4.07*** | | ***10.00*** | | ***17.97*** | | | ***8.32*** | | ***0.75*** | | ***0.42*** | |  |  |  |  |

| Table S10. Detailed Pilot Valley Geochemistry - Hydration and Carbon by Group  BDL=Below Detection Limit ND=No Data | | | | |
| --- | --- | --- | --- | --- |
| Core ID | TOC | TIC | Total Carbon | Avg. Hydration |
|  | wt% | wt% | wt% | wt% |
| **Group 1** | | | | |
| PV2-2 | 0.54 | 2.63 | 3.17 | 21.80 |
| PV2-3 | 1.10 | 1.98 | 3.08 | 25.40 |
| PV2-4 | 0.72 | 2.40 | 3.12 | 26.20 |
| PV2-5 | 0.36 | 1.58 | 1.94 | 26.60 |
| PV5-4 | 0.75 | 1.67 | 2.42 | 27.0 |
| ***Average*** | ***0.69*** | ***2.05*** | ***2.75*** | ***25.40*** |
| **Group 2** | | | | |
| PV2-6 | 0.88 | 1.94 | 2.82 | 27.70 |
| PV2-7 | 0.42 | 1.01 | 1.43 | 29.20 |
| PV2-8 | 1.40 | 5.01 | 6.41 | 25.10 |
| PV2-9 | 0.26 | 4.07 | 4.32 | 28.10 |
| PV3-2 | 0.78 | 1.45 | 2.24 | 23.7 |
| PV3-3 | 0.16 | 0.34 | 0.50 | 23.80 |
| ***Average*** | ***0.65*** | ***2.30*** | ***2.95*** | ***26.27*** |
| **Group 3** | | | | |
| PV1-4 | 0.28 | 3.25 | 3.52 | 26.60 |
| PV1-5 | 0.42 | 3.85 | 4.27 | 31.60 |
| PV2-1 | 0.43 | 1.70 | 2.13 | 31.60 |
| PV2-10 | 0.70 | 5.79 | 6.49 | 31.00 |
| PV2-11 | 0.96 | 5.07 | 6.03 | 38.50 |
| PV3-1 | 0.88 | 0.97 | 1.9 | 27.0 |
| PV5-1 | ND | ND | ND | ND |
| PV5-2 | 0.13 | 0.28 | 0.40 | 24.1 |
| PV5-3 | 0.24 | 0.01 | 0.25 | 24.2 |
| ***Average*** | ***0.51*** | ***2.62*** | ***3.12*** | ***29.33*** |

**Supplementary Figures**

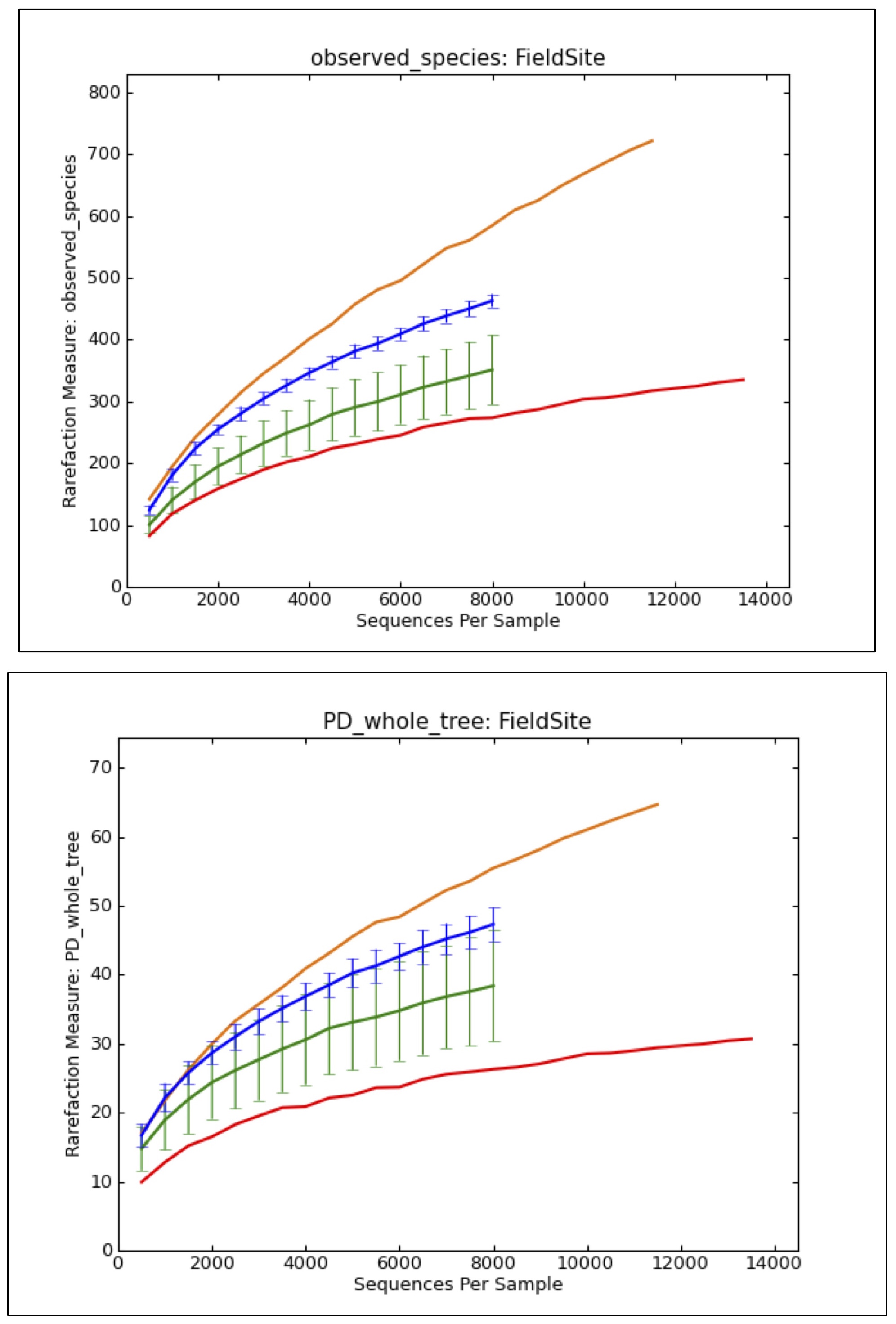

Figure S1. Rarefaction plots for field site comparison study. a) Observed species. b) Faith's phylogenetic diversity [*Faith and Baker*, 2006].

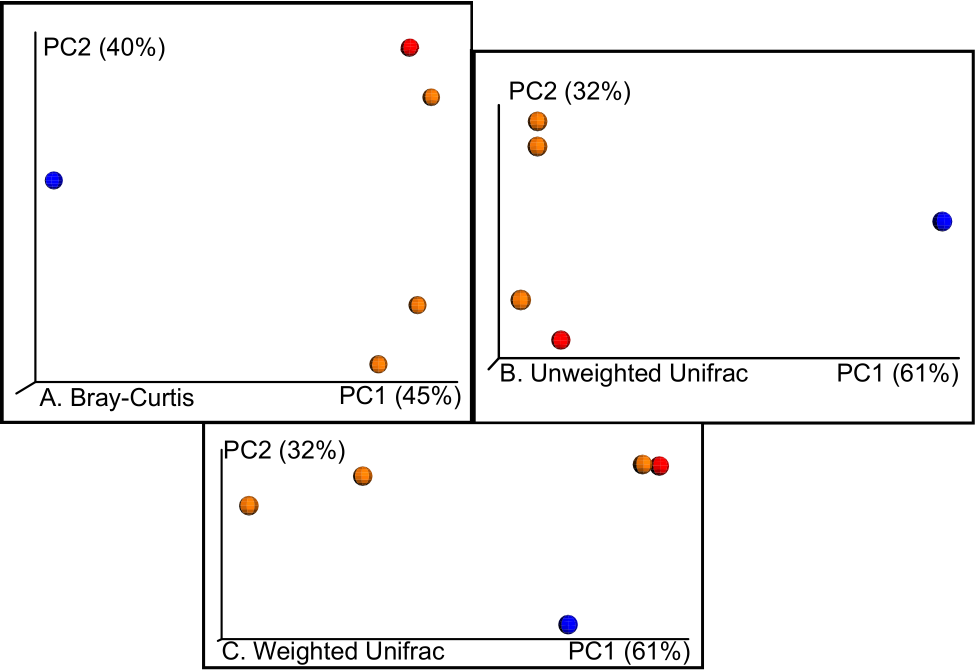

Figure S2. PCoA plots of a) Bray-Curtis, b) un-weighted Unifrac, and c) weighted Unifrac beta diversity metrics. Blue = Pilot Valley, Red = Bonneville Salt Flats, Orange = Great Salt Lake.

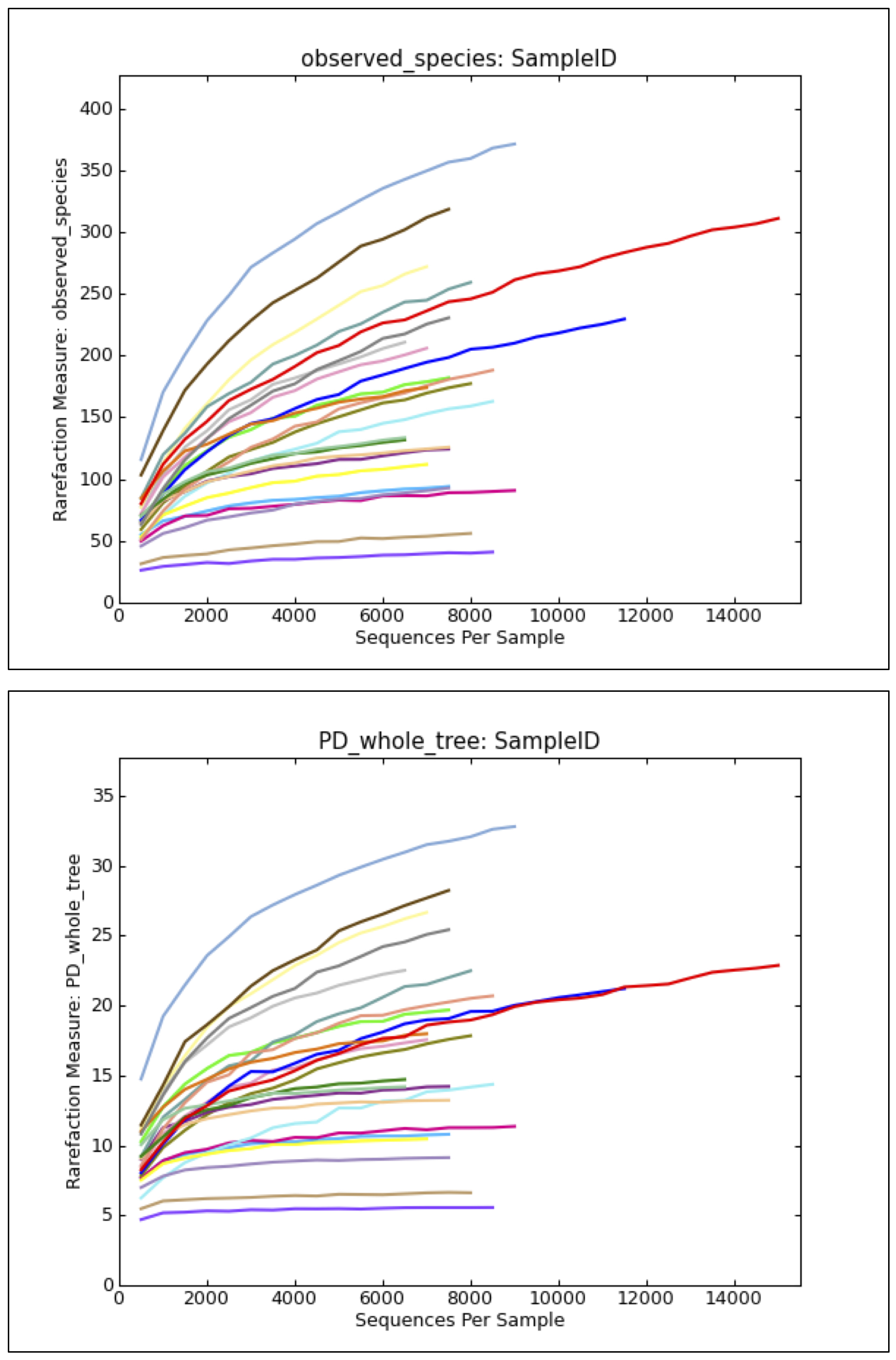

Figure S3. Rarefaction plot of sample coverage for transect study. Each rarefaction curve represents each individual transect sample per Figure 1. top) observed species; bottom) Faith’s phylogenetic diversity [*Faith and Baker*, 2006].

**SI References**

Faith, D. P., and A. M. Baker (2006), Phylogenetic diversity (PD) and biodiversity conservation: some bioinformatics challenges, *Evolutionary Bioinformatics Online*, *2*, 121-128.
